# Multi-context seeds enable fast and high-accuracy read mapping

**DOI:** 10.1101/2024.10.29.620855

**Authors:** Ivan Tolstoganov, Marcel Martin, Nicolas Buchin, Kristoffer Sahlin

**Affiliations:** Department of Mathematics, Science for Life Laboratory, Stockholm University, 106 91, Stockholm, Sweden; Department of Biochemistry and Biophysics, National Bioinformatics Infrastructure Sweden, Science for Life Laboratory, Stockholm University, Box 1031, SE-17121 Solna, Sweden

**Keywords:** read mapping, seeds, Illumina, k-mers, strobemers

## Abstract

A key step in sequence similarity search is to identify shared seeds between a query and a reference sequence. A well-known tradeoff is that longer seeds offer fast searches but reduce sensitivity in variable regions. We introduce multi-context seeds (MCS), which allow to store seeds with different lengths in the same index structure, thus retaining the advantages of both short and long seeds. We demonstrate the applicability of MCS by implementing them in strobealign. Strobealign with MCS substantially improves accuracy compared to the previous version with little cost in runtime and no memory overhead.

## Background

In bioinformatics, a *seed* refers to a short substring or subsequence of a biological sequence that is used to initiate a sequence similarity search between sequences, e. g., between a read and a reference genome. Seeds are crucial for heuristic (i. e., not guaranteed to find the optimal solution) algorithms for sequence comparison such as read mapping algorithms [1, 38]. In the seeding step, the seeds help to quickly identify potential *matches*, reducing the search space. A match (also sometimes referred to as an anchor) is a seed that occurs both in the query and the reference.

A *k*-mer is a string of length *k* and is commonly used in sequence similarity search algorithms such as read mapping. In read mapping, choosing *k* is a tradeoff: Longer *k*-mers enhance seed specificity and therefore mapping speed, but shorter *k*-mers increase sensitivity as they are less affected by sequencing errors and biological mutations. Several alternative methods have been proposed to enable using longer seeds that can match in the regions between mismatches and indels. These include spaced seeds [28], TensorSketching [17], context-aware seeds [44], *k*-min-mers [9], strobemers [36], BLEND [12], and SubSeqHash [27, 26]. Some of these seed types were shown to be efficient for short [37] and long-read mapping [10, 12, 17] and can enable one or two orders of magnitude speedup over using *k*-mers [10, 37, 12, 17]. At a high level, the seeds in [10, 37, 12] are based on joining together several *k*-mers into a single seed, thus providing a longer range of a match, which gives more unique seeds and, thus, fewer candidate locations to explore. However, long seeds, although they allow approximate matches, come with an undesirable reduction in accuracy, which results in widespread usage of the more resource-intensive but accurate short seed-based alternatives [22, 24, 25, 40].

The tradeoff between seed length and accuracy makes the idea of having seeds at multiple levels or resolutions compelling. The longer seeds would provide speed through their low match redundancy, while one could fall back on shorter seeds in case no longer matches were found. The concept of using different *k*-mer sizes for various resolutions has been used in, e. g., genome assembly [33, 3, 2], transcriptome assembly [35], RNA-Seq analysis [19], metagenomic mapping [41] through LexicHash [14] and *k*-mer counting [18]. In read mapping, variable-length seeds such as MEMs [6, 24] (constructed with the FM-index [11] or Enumerated Radix Tree [42]) and MCAS [16] are used, but they are slow to query. This is because the query process to construct such seeds needs to access scattered locations in memory. Moreover, aforementioned multi-resolution methods only increase or decrease the resolution with one nucleotide [24] or utilize multiple sizes of *k* [3], which results in a runtime increase proportional to the number of *k* sizes used.

## Results

### Method overview

A *strobemer* is a seed that consists of two or more *k*-mers called *strobes*. We propose using strobemers that support queries at multiple resolutions, making it possible to query all or a certain subset of these *k*-mers in the index. Naively, we could store both *k*-mers and strobemers with 2, … *n k*-mers in the same index. However, this would increase space requirements and would also slow down mapping through cache misses if the seeds are scattered in memory. Instead, we present *multi-context seeds* (MCS). MCS are strobemers produced with the randstrobe method [36] that distribute the bits allocated to store the (hash of the) seed separately between *k*-mers in the strobemer. The primary idea of MCS is as follows. For an *L* bits allocation of a seed hash value, an MCS will store the leftmost *l*_1_ *< L* bits to represent the first *k*-mer in the strobemer, the following *l*_2_ *< L* bits for the second *k*-mer, etc., with *l*_1_ + *l*_2_ + … = *L*. See Fig. 1 for a conceptual illustration. When indexed efficiently, for example as a vector sorted by hash values paired with a look-up table, as is standard in *k*-mer based mapping [25, 46, 37], this seed enables cache-efficient lookup for both *full matches* covering the whole strobemer and *partial matches* covering the first several strobes.

**Fig. 1.**
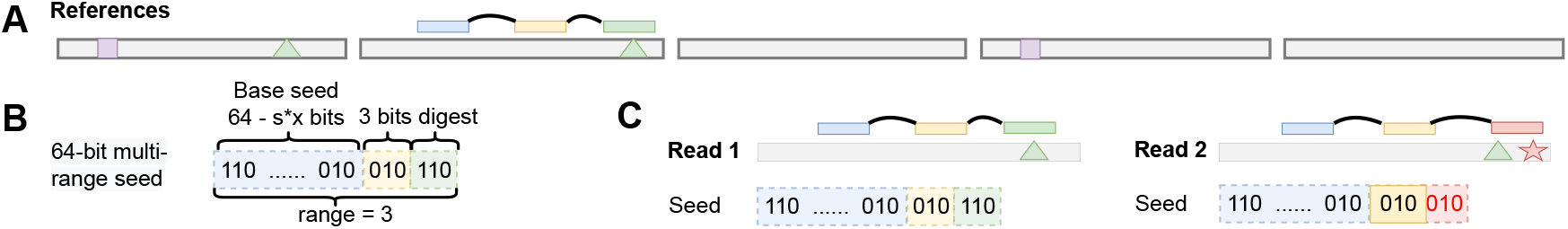
Multi-context seeds concept. **A** A reference sequence consisting of five similar copies with some variation (colored shapes), and a strobemer seed sampled from one of the copies. **B** The forward multi-context seed representation of the strobemer consisting of a (64 − *sx*)-bit prefix for the base strobe, and *x*-bit prefixes for the *s* remaining strobes. In this example, *s* = 2 and *x* = 3. **C** Read mapping of a read without sequencing errors (read 1) and a read with a sequencing error or non-reference mutation in the third strobe (read 2). While read 2 cannot produce a full match, the part of the seed limited to the first two strobes is still able to produce a match to the reference.

Being able to search for seed matches at several resolutions compared to the binary “match or no match” of the full seed increases both accuracy and percentage of aligned reads. We first demonstrate that MCS improve sequence matching characteristics over randstrobes and *k*-mers but retain roughly the same seed uniqueness benefits as randstrobes. We then show the practical applicability of MCS by implementing them in strobealign [37] in addition to randstrobes. We benchmark strobealign with MCS and compare it to strobealign with randstrobes and other state-of-the-art alignment tools using simulated Illumina reads of different lengths from four genomes at various sizes and sequence complexities. Strobealign with MCS improves mapping-only and extension-based read alignment accuracy over strobealign with randstrobes in all experiments with typical Illumina read lengths ≤ 150 nt for both single-end and paired-end data, without a substantial slowdown. Particularly, as sequence variability gets high, MCS have a higher relative increase in accuracy over strobealign with randstrobes. Unlike strobealign with randstrobes, strobealign with MCS now surpasses minimap2 accuracy in extension alignment mode in nearly all experiments, while remaining substantially faster. When compared with the gold standard BWA-MEM, strobealign with MCS has comparable accuracy on datasets with high diversity, but with much lower mapping and indexing time.

### Evaluation overview

We first compare multi-context seeds with randstrobes and *k*-mers, taking into account sequence matching coverage to assess sensitivity and seed uniqueness to assess specificity. This is a general-purpose sequence similarity search benchmark to assess the repetitiveness of seeds and match sensitivity in a general setting. We then implement multi-context seeds in addition to randstrobes in the read mapping tool strobealign [37] (published as v0.17.0). We benchmark the multi-context seeds version against the randstrobe version of strobealign, BWA-MEM (v0.7.19) [24], and minimap2 (v2.30) [25] using simulated and biological short-read data. We compare against BWA-MEM and minimap2 as they were the competitors with the highest accuracy (BWA-MEM) and most desirable speed-to-accuracy tradeoff (minimap2) to strobealign in a previous benchmark [37] that included other tools such as Bowtie2 [22], SNAP2 [5], and Accel-Align [46]. Additionally, we report benchmark results for the recently released X-Mapper tool [13] in the supplement (see Suppl. Section 4 for details). We also benchmark MCS against randstrobes and a *k*-mer only version of strobealign as well as minimap2 in a proof-of-concept long-read mapping benchmark. Support for long reads is in development and only available in mapping-only mode in strobealign v0.17.0 (i. e., without base-level alignments).

### Datasets

For the general-purpose sequence matching benchmark, we simulate pairs of nucleotide sequences of length 10,000 with mutation rates 0.01, 0.05, and 0.1 (insertions, deletions, or substitutions occurring with equal probability 1*/*3). Each sequence pair is replicated 1000 times to reduce sample variability. We refer to these pairs of simulated sequences as the **SIM-R** dataset. Such experiment setup was performed in [36] to evaluate different strobemer constructions. For the uniqueness evaluation, we measure the fraction of unique seeds (occurring once) and the E-hits [37] in the human cell line CHM13 assembly [31], referred to as **CHM13**.

For the read alignment benchmark, we generate three datasets with different mutation rates that we call **SIM0, SIM4**, and **SIM6. SIM4** has already been used in an earlier study [37].

Simulated reads were generated from the CHM13 (3.1 Gbp), fruit fly (144 Mbp), maize (2.2 Gbp) and rye [23] (7.3 Gbp) genomes. Libraries of paired-end reads of various lengths ranging from 50–500 nt are generated, and single-end libraries are obtained by omitting the second reads of paired-end reads. We also simulate single-end reads of lengths 1000, 5000, and 10,000 nt (long reads).

**SIM0** consists of reads without errors and without variants, i. e., the reads are substrings of the reference. **SIM4** and **SIM6** are generated with Mason [15] simulating Illumina reads at different mutation rates. Variants simulated for **SIM4** occur at a rate somewhat higher than expected between humans. The rate is much higher for **SIM6** (see Suppl. Section 5 for details). Note that even the long reads were generated using the Illumina read profile (using Mason) as a way to generate PacBio-HiFi-like reads (see Suppl. Section 5).

We also performed a SNV and indel calling benchmark as performed in [37], using bcftools as the variant caller with two simulated (**SIM150, SIM250**) and two biological (**BIO150, BIO250**) datasets.

### Sequence matching experiment

#### Benchmark

We perform the sequence matching benchmark of seeds similarly to the procedure described in [36]. Given a pair of simulated sequences (*s, s*^*′*^) from the **SIM-R** dataset, we construct *k*-mers, randstrobes, and MCS from every possible starting position of the two sequences. We count the number of matches between seeds generated from *s* and seeds generated from *s*^*′*^ (including partial matches for MCS) for every type of seed construct. Similarly to the strobemers study [36], we compute match percentage, sequence coverage, and the island E-size metric as defined in [36]. Briefly, match percentage *m* is the fraction of seed matches, sequence coverage *sc* is the percentage of sequence in *s* covered by the actual strobes of a seed. The island E-size is a measure of the size of gaps (islands [4]) without matches. We describe these metrics in detail in Supplementary Section 3. An additional match coverage metric, which is a percentage of sequence spanned by seed matches, is included in Suppl. Tab. S1, with results similar to sequence coverage. As expected, the multi-context seeds result in the same or higher number of matches and sequence coverage compared to *k*-mers with the same *k* parameter and randstrobes with the same parametrization, with the exception of multi-context seeds with (2, 15) parametrization, which result in slightly lower sequence coverage metric than 15-mers. This discrepancy can be explained by the fact that randstrobes are not sampled if the distance from the start position to the end of the sequence does not exceed 2*k* + *w*_min_. Therefore, *k*-mer matches located close to the end of the sequence will not have corresponding partial MCS matches.

#### Results

Table 1 shows the mean match percentage, total sequence coverage, and island E-size for the **SIM-R** dataset. We benchmarked randstrobes and MCS with parametrizations (2, 15) and (3, 10) with total strobemer length 30 (as used in [36]). For the *k*-mers, we chose *k* size 30 corresponding to the total strobemer size, and *k* sizes 10, 15, and 20 corresponding to all the possible partial match sizes of the multi-context seed parametrizations. As expected, we observe that *k*-mers of size 10 and MCS with parametrization (3, 10) have the highest fraction of matches, since the seeds of smaller size are more likely to be preserved between mutation positions. Between the seeds of the same length, i. e. randstrobes, MCS, and *k*-mers of size 30, MCS with parametrization (3, 10) show the highest match percentage due to production of both full and partial matches. MCS have higher match coverage and sequence coverage than randstrobes due to the presence of additional partial matches.

**Table 1.**
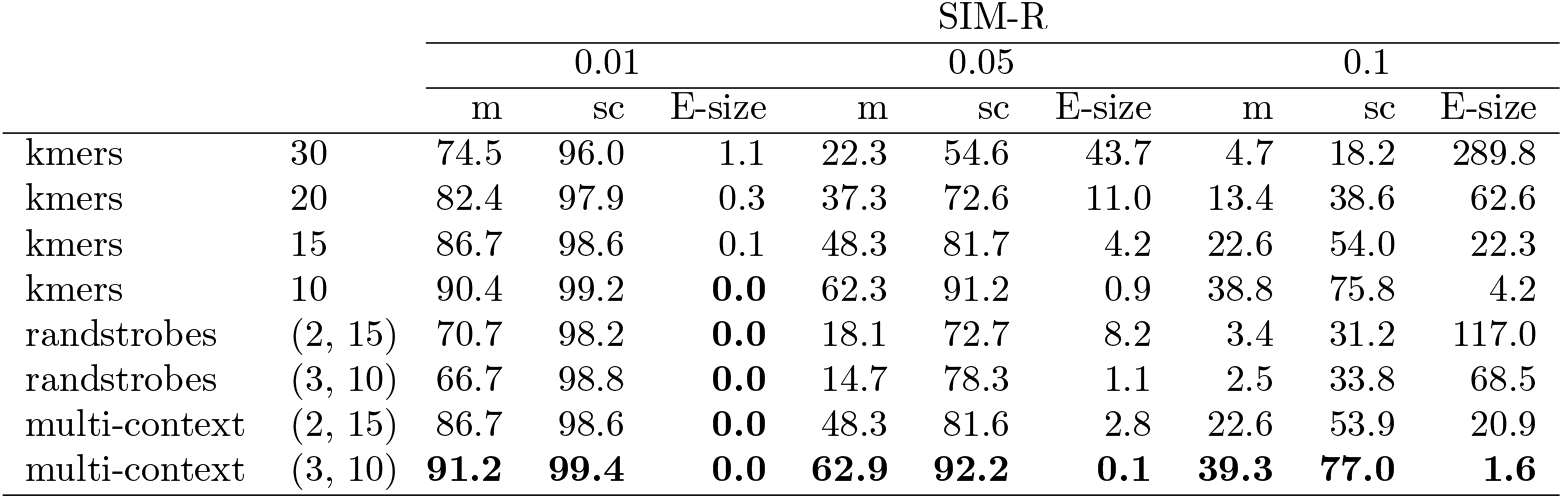
Match statistics for *k*-mers, randstrobes, and MCS under mutation rates of 0.01, 0.05, and 0.1. Here, *m* denotes the number of matches (both full and partial for MCS) as a percentage of the total number of extracted subsequences for the protocol, *sc* (sequence coverage) is shown as the percentage of the total sequence length, and *E*-*size* is the expected island size. The randstrobes and multi-context seeds were constructed with the window parameters *w*_min_ = 25 and *w*_max_ = 50

MCS demonstrate significant advantage over randstrobes with the same parameters. For example, for the parametrization (3, 10) and mutation rate 0.1, the match percentage is 39.3% for MCS and 2.5% for randstrobes, sequence coverage is 77% for MCS and 33.8% for randstrobes, and the expected island size is only 1.6 for MCS and 68.5 for randstrobes. Also, the improvement of MCS over randstrobes for the two-strobe setting (2, 15) is almost as large as for the three-strobe setting.

### Seed uniqueness

#### Benchmark

While MCS show higher match coverage than randstrobes and *k*-mers due to the presence of partial matches, they are also expected to result in a larger amount of spurious matches for the same reason. The previous experiment was designed to be without repeats to accurately measure (true) coverage of matches. In order to assess the potential increase in the spurious matches, we benchmarked MCS against *k*-mers and randstrobes on the CHM13 reference genome using three metrics, namely the percentage of unique seeds, number of distinct seeds, and the E-hits metric [37]. Roughly, E-hits measures the expected number of occurrences of a seed if a seed is selected uniformly at random in the database, see [37] for details. We measure the seed uniqueness metrics by indexing the CHM13 reference genome and counting the seed occurrence distribution. We estimate the *k*-mer occurrence distribution using KMC [21] *k*-mer counter tool, and strobealign for calculating randstrobes and MCS occurrence distribution. For this experiment, we use all *k*-mers in the reference in strobealign, i. e., we do not use the open-syncmers downsampling in strobealign. For the MCS, we provide the number of partial hits separately from the number of full hits. We use a range of typical *k*-mer sizes used for human genome applications and resulting matching strobemer parameterizations (dividing the *k*-mer size by 2 or 3). We used 56 bit hashes (the number of bits allocated to a seed hash value in strobealign) for randstrobes and multi-context seeds. In MCS, 40 bits were reserved for the first hash, and 16 bits for the second (in the case of 2-strobe MCS) or 8 bits for the second and for the third (in the case of 3-strobe MCS).

#### Results

We observe that MCS (full hits) and randstrobes are nearly identical in uniqueness (Table 2). This suggests that even though MCS splits the bits in the hash value representation to enable multi-level search, it does not result in significant hash collisions for full matches compared to randstrobes, at least not in a human genome. As expected, partial matches are more redundant than full matches. Also, the partial hits of the MCS show similar proportion of unique seeds and higher E-hits to *k*-mers of the same length, and in the three-strobe setting, the partial hit with first two strobes is less unique than k-mers of the same length 32. This difference between *k*-mers and partial hits can be attributed to the hash collisions, since the second strobe is encoded in 8-bit hash in this benchmark.

**Table 2.**
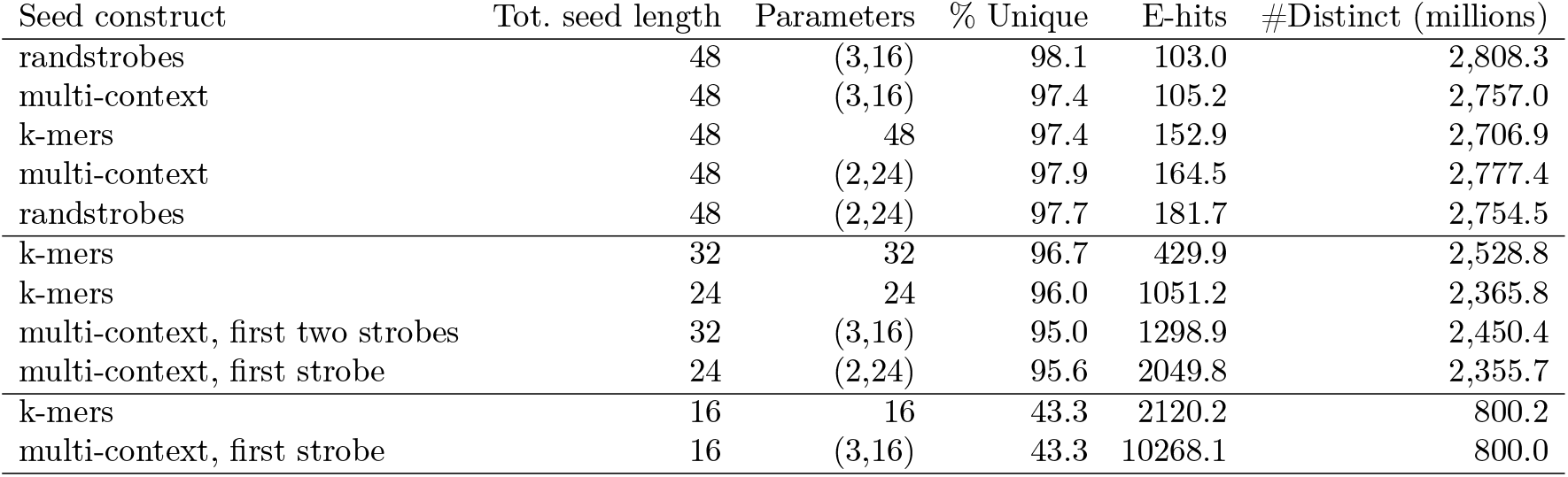
Uniqueness statistics for *k*-mers, randstrobes, and MCS. For the MCS, every given seed is estimated separately as its full hash value, subhash corresponding to the first strobe, and subhash corresponding to the first two strobes for 3rd order MCS. The column “Tot. seed length” indicates total nucleotides included in the seed. “% Unique” denotes the percentage of the total number of seeds of the given type that have unique hash value, “E-hits” denotes the E-hits metric of the given seed type, and “#Distinct” shows the number of distinct seeds of the given type (in millions). The rows are sorted by the E-hits value. The randstrobes and multi-context seeds were constructed with the window parameters *w*_min_ = 36 and *w*_max_ = 72. For both the randstrobes and multi-context seeds, we used 56-bit hashes, with 32 bits reserved for the MCS base hash.

### Read mapping evaluation

We compared strobealign-MCS to strobealign with randstrobes, BWA-MEM and minimap2. For the single-end and paired-end reads experiments, we show the accuracy and runtime for the **SIM0, SIM4** and **SIM6** datasets for CHM13 in Fig. 2 and Fig. 3. As in [37], we consider a read correctly mapped if the reported mapping coordinates overlap with the provided ground truth reference coordinates for the read. Otherwise, it is incorrectly mapped. Accuracy is computed as the fraction of correctly mapped reads. Results for all four genomes (also rye, fruit fly, and maize) and measurements of memory usage and mapping rate (percentage of aligned reads) are found in Suppl. Figs. S8–S14.

**Fig. 2.**
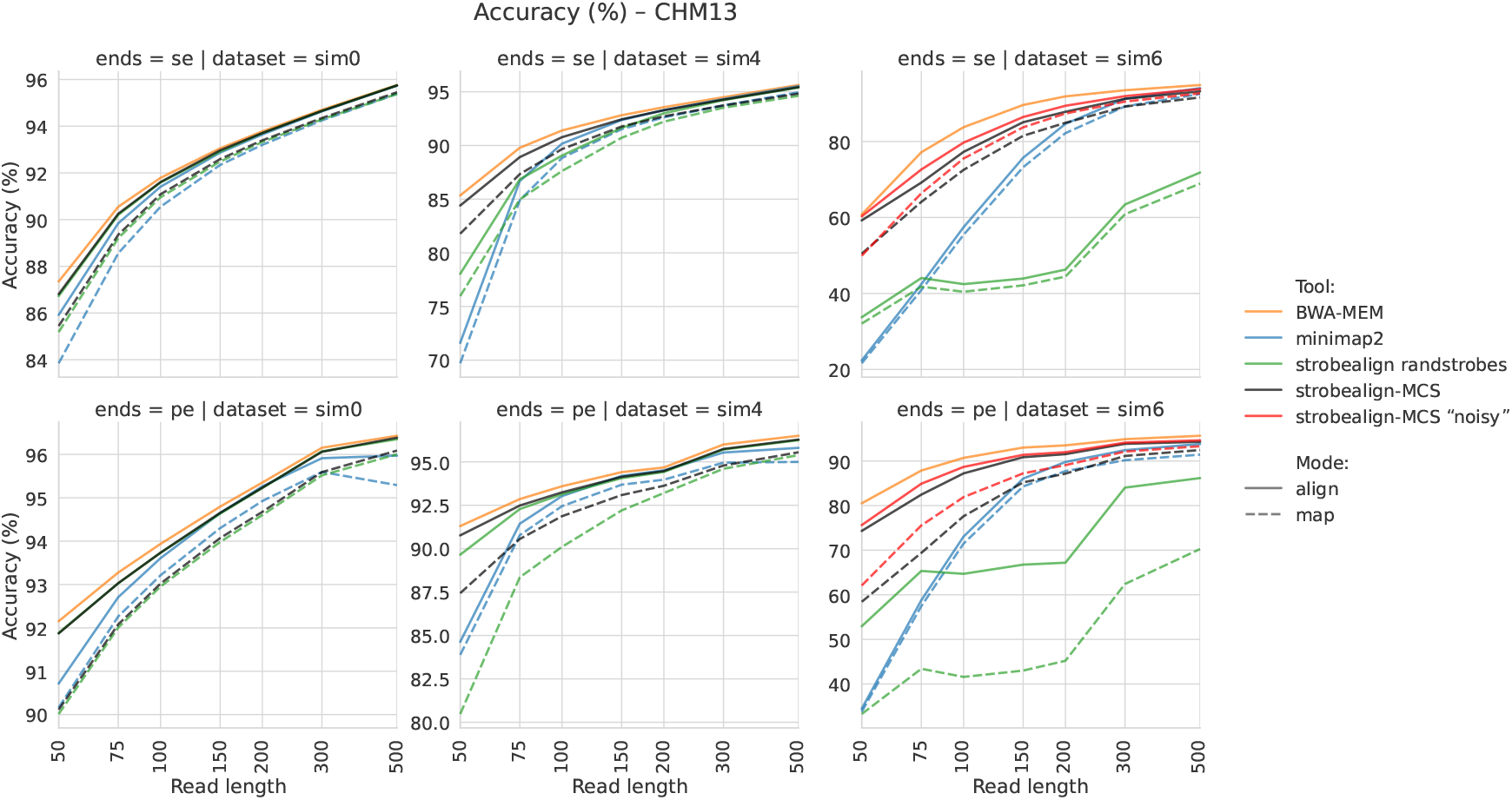
Accuracy of single-end (top row) and paired-end (bottom row) read alignment for the **SIM0** (left), **SIM4** (middle), and **SIM6** (right) datasets. Mode “align” is extension-alignment mode (SAM output), mode “map” is mapping-only mode (coordinate output). The *x*-axis (read length) is scaled logarithmically.

**Fig. 3.**
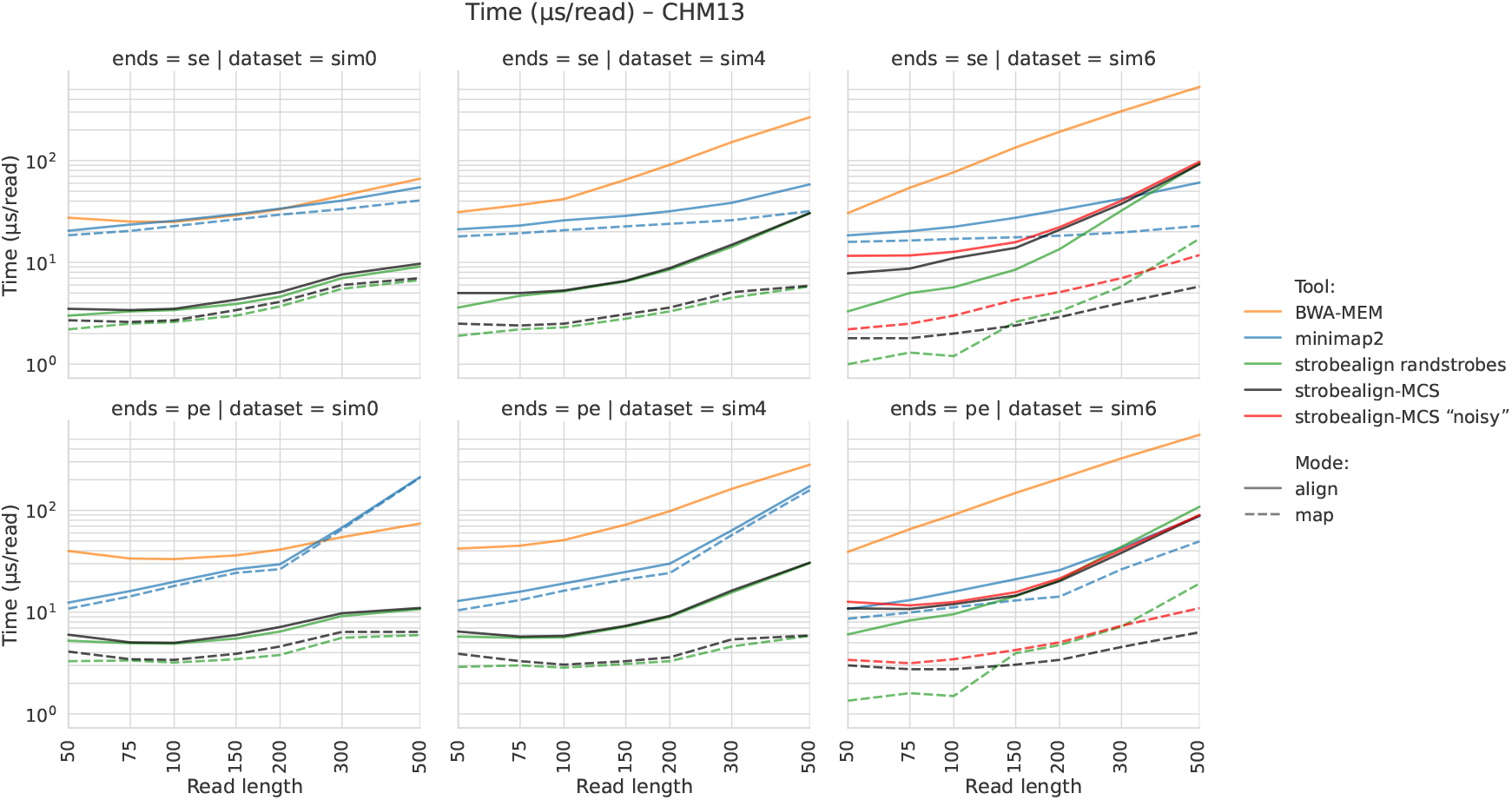
Runtime of single-end (top row) and paired-end (bottom row) read alignment for the **SIM0** (left), **SIM4** (middle), and **SIM6** (right) datasets

#### High-variation profile

Similarly to minimap2’s several mapping profiles, we implemented two profiles, a high-quality profile (same parameters as strobealign for short-reads) and a high-variation profile that we label *noisy*. The noisy profile sets *k* = 16, *s* = 2, *w*_*min*_=2, *w*_*max*_=2, which, in practice, produces 2-layer *k*-mers, where the smaller *k*-mer is 16 and the larger is 26 (in expectation; depending on downstream syncmer offset).

#### Short-read mapping results

For paired-end reads, strobealign-MCS improves both accuracy (Figure 2, Suppl. Fig. S9) and the percentage of aligned reads (Suppl. Fig. S13) over strobealign with randstrobes, both in mapping-only and extension-alignment modes in all genomes. The improvement of MCS over randstrobe seeds in the alignment mode is present for all datasets, and is most apparent for reads up to length 100 bp (mean accuracy increase 0.7%) in the **SIM4** dataset (moderate mutation rate), and for all read lengths in the **SIM6** dataset (mean increase 17.2%). This is expected as the partial seed matches can produce hits around variations and other divergent regions where no full matches or randstrobes were produced. In the mapping-only mode (no extension alignment), the increase in accuracy is noticeable (mean 2.2%) for read lengths 300 bp or lower in the **SIM4** dataset and for all read lengths in the **SIM6** dataset (mean 28.2%). The mapping-only mode is useful for rapid metagenomics analysis with the AEMB mode [32]. In both read-mapping and extension modes, strobealign with MCS further narrows the mean accuracy gap for shorter reads compared to gold-standard BWA-MEM, while still being 6× faster on average in the alignment mode (Suppl. Fig. S11).

For single-end reads, the relative increase in accuracy of strobealign-MCS compared to strobealign with randstrobes is even larger, visible for reads shorter than 200 bp in the alignment mode (Figure 2, Suppl. Fig. S8). Relatively better performance in the single-end mode can be explained by strobealign’s rescue procedure in paired-end mode, which utilizes the matches of a read to recover its unmapped mate. In addition, strobealign with MCS also substantially improves the percentage of mapped reads over strobealign with randstrobes for shorter read lengths (Suppl. Fig. S12).

For SIM0 and SIM4 datasets, the runtime overhead of strobealign-MCS to strobealign with randstrobes is minimal, with substantial accuracy gains for shorter read lengths. For SIM6, strobealign-MCS is slightly slower than strobealign with randstrobes (Figure 3, Suppl. Figs. S10 and S11), due to the higher redundancy and extra lookups of partial hits, as well as the extra alignment sites produced by partial matches found by strobealign in the absence of full matches.

#### Proof-of-concept long-read mapping results

To assess MCS outside of short-read mapping, we implemented a proof-of-concept long-read mapping mode in strobealign by adding a seed chaining algorithm (described in Suppl. Section 8). We consider it proof-of-concept because this mode currently only works in mapping-only mode (no base-level alignments), nor are supplementary mappings implemented (needed for, e. g., structural variation detection with long reads). We simulated long-reads (read lengths 1,000, 5,000, and 10,000 nt, Suppl. Section 5.3) for long-read mapping.

Results are shown in Suppl. Figs. S6 and S7. Firstly, strobealign-MCS substantially outperform strobealign with randstrobes, particularly in modest to high variation settings, while also being faster in most cases.

Additionally, compared to state-of-the-art long-read mapper minimap2, strobealign with MCS is 3-5× and 1.2-2.3× faster with both default and noisy profiles, respectively. Additionally, the two profiles are also often more accurate than minimap2. For example, on CHM13 with no or modest variation (SIM0 and SIM4) strobealign has higher accuracy than minimap2. Overall, across the 27 experiments (genomes, variation levels, and read lengths), strobealign demonstrates comparable accuracy levels to minimap2, while improving runtime.

#### Strobealign seed lookup strategies

With the implementation of MCS in strobealign, it can now use different seeding approaches; randstrobes-only, *k*-mer-only (only searching for the base hash corresponding to a single syncmer; see methods), and MCS. We compared these settings in strobealign (Suppl. Fig. S5). Overall, we see that MCS produce favorable accuracy over randstrobes mode while producing comparable results to *k*-mer-only. However, for larger and more repetitive datasets, MCS is faster than *k*-mer-only.

#### 3-strobe based approach

While the current version of strobealign is based on MCS consisting of two strobes, we developed an additional 3-strobe based version to assess how increasing the number of search layers would affect accuracy and runtime (see Suppl. Sec. 7 for details). Within strobealign’s mapping framework, our 3-strobe implementation did not result in any additional benefits compared to 2-strobe version (Suppl. Figs. S15 and S16).

#### Indexing time

Indexing speed is not measurably affected by enabling the MCS hash function. Using four threads and leaving all other indexing parameters (*k, s*, etc.) unchanged, strobealign takes 2 s, 32 s, 47 s and 111 s to index fruit fly, maize, CHM13, and rye, respectively. For reference, BWA and minimap2 take 2420 s and 52 s, respectively, to index CHM13 on the same machine (using four threads for minimap2; BWA does not parallelize index creation).

#### Variant calling

We ran the variant calling benchmark on two simulated and two biological Illumina datasets as performed in [37] (Suppl. Fig. S17). In terms of F-score, the tools perform roughly similar. BWA-MEM has highest SNV recall for all datasets, while the tools perform similarly on indels. Note however that we use bcftools to call variants. Some callers may be developed or tuned based on popular aligners’ mapping quality (MAPQ) scores. Specifically, it has been shown that *bcftools call* was the variant caller that produced the best results with BWA-MEM alignments out of seven variant calling tools [47]. Therefore, we view this analysis as a sanity check rather than a perfect indicator of mapping quality. As with the simulated data, strobealign is substantially faster than minimap2 and BWA-MEM. There is no significant difference between strobealign versions in calls. While MCS enables looking for partial matches and would enable split or supplementary mapping, it is not yet implemented in strobealign. We hope to include split read mapping in future work, which may improve variant calling further.

## Methods

### Preliminaries

#### Notation

Let *R* be a string over the alphabet *Σ* = {*A, C, G, T* }. We refer to a string obtained by deleting a subset of characters in *R* as a *subsequence* of *R*. We define the consecutive subsequence as a *substring*. We will use zero-indexed notation for strings. Given nucleotide string *R*, a *k-mer* is defined as a substring of *R* of length *k*. Let *R* [*i, k*] denote a *k-mer* covering the positions from *i* to *i* + *k* − 1. We use *h*_*L*_ to denote a hash function that maps *k*-mers to *L*-bit strings *h*_*L*_ : *Σ*^*k*^ → {0, 1}^*L*^. The *l*-bit prefix of the *L*-bit hash *h*_*L*_ is denoted as 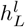.

#### Strobemers

Strobemers were introduced in [36] as a way of handling indels and reads with a high mutation rate. Strobemers are constructed from an array *K* = (*s*_1_, …, *s*_*N*_) obtained by decomposing a string *R* into *N* consecutive *k*-mers, where *s*_*i*_ = *R*[*i, k*]. Given *K* and parameters (*n, k, w*_min_, *w*_max_), a *strobemer* of order *n* is defined as a subsequence of *n k*-mers 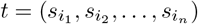 in *K*. The *w*_min_ and *w*_max_ parameters limit the distance between consecutive *k*-mers in *t*, i. e. for every *j* ∈ [1, *n* − 1], we have *i*_*j*+1_ ∈ [*i*_*j*_ + *w*_min_, *i*_*j*_ + *w*_max_]. The *k*-mers selected for a strobemer are called *strobes*.

Strobemers as initially described were constructed from complete *k*-mer arrays *K* of a reference. However, strobemers can also be constructed from a subset of *k*-mers. For example, in strobealign [37], strobemers are constructed from the array of *open syncmers* [8] generated in the reference. In this work, we abstract over the array of *k*-mers from which a strobe is chosen by denoting 𝒲_*j*_ to be the indices of *k*-mers available in a window where strobe *j* is selected. When selecting strobes from all *k*-mers, the window corresponds to the consecutive indices of a substring of the reference of length *w*_max_ − *w*_min_ + 1, while it is a subsequence of reference indices with larger span if selecting over a subset of *k*-mers such as minimizers [39, 34, 30] or syncmers [8].

#### Randstrobes

Various strategies for selecting the next strobe were described in [36, 29]. Randstrobes are constructed iteratively from the starting strobe 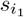. If strobes 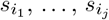 were already selected for a randstrobe, the strobe 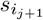 is selected as

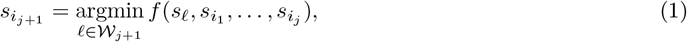

where *f* is a function that maps a strobe to an integer. Several functions were investigated in [20]. For example, strobealign uses

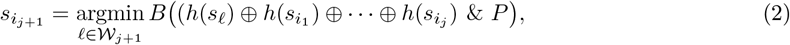

where *h* is a hash function from a sequence to a 64-bit number, *B* is the number of set bits, *P* is a bitmask with its *p* least significant bits set, and ⊕ is the bitwise XOR operator. The *B* and *p* make strobealign pick strobes earlier in the window more often, which is deemed suitable for the shortest reads (for details, see [37]). To combine the hashes of individual strobes into a single hash value for the randstrobe, strobealign v0.15.0 and earlier uses (*h*(*a, b*) = *h*(*b, a*)) hash function 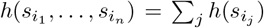 to hash strobes from two different orientations into the same hash value, as described in [37]. Since v0.16.0 strobealign uses multicontext seed hash function (Eq. 3) to construct a randstrobe hash value in order to enable randstrobes and multi-context seeds within the same index.

### Multi-context seeds

We introduce multi-context seeds, a seeding strategy based on the strobemers [36] technique. Strobemers consist of several *k*-mers, or *strobes*, such that the distance between two consecutive *k*-mers does not exceed a constant parameter. While this strategy allows to effectively increase the length of the seed, it results in lower sensitivity, especially in shorter reads. To alleviate this, we change the strobemer hash function to a concatenation of prefixes of individual strobe hashes. This hash function allows searching for the first consecutive strobes instead of the whole strobemer by using a prefix of the hash in a search, as illustrated in Figure 1. Effectively, this means that for strobe length *s*, seeds of lengths *s*, 2*s*, …, *ns* are stored simultaneously in a single index. A formal description of multi-context seeds and their implementation in strobealign [37] follows.

#### Multi-context seeds

Given integers *L, n*, and *l*_1_, …, *l*_*n*_ such that 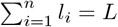, the *multi-context* seed is a strobemer consisting of strobes 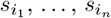, which is stored as

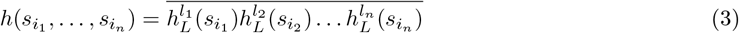

where 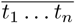 denotes the concatenation of bit strings *t*_1_, …, *t*_*n*_.

For example

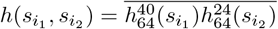

describes a strobemer with two strobes hashed in a 64-bit integer where the first strobe is represented with 40 bits and the second strobe with 24 bits. For an illustration, see Fig. 1. We say that the first *l*_1_ bits of the MCS hash are the *base part* of the hash, and the following allocations are (shorter) *digests*. Typically, these are given more bits than downstream strobes to avoid hash collisions of the base *k*-mer.

### Implementation in strobealign

#### Open syncmers

Open syncmers is a *k*-mer subsampling method [8]. Given positive integer parameters (*k, s, t*), where *s* ≤ *k* and *t* ∈ [0, *k* − *s* + 1], a *k*-mer is selected as an *open syncmer* iff the smallest of its *k* − *s* + 1 consecutive *s*-mers occurs at position *t*. The smallest *s*-mer is defined based on a hash function. In order to process queries from both the forward and the reverse strands efficiently, we store a canonical representation of the open syncmers, i. e., only the smallest syncmer out of the forward and the reverse-complement *s*-mers.

#### Multi-context seeds in strobealign

As in the randstrobe version of strobealign, we use the canonical open syncmers in order to create the sequence of strobes for multi-context seed construction. We construct open syncmers using the parameters *k, s*, and *t* = ⌈(*k* − *s* + 1)*/*2⌉, where parameters *k* and *s* are dependent on the read length (see details in the Supplementary Section 1). Depending on the value of *t*, 14 − 20% of *k*-mers are selected as strobes. Following the construction of the sequence of open syncmers *S* = (*s*_1_, …, *s*_*N*_) from the input references, we employ the *skewed randstrobe* selection strategy described in strobealign [37] for both randstrobe and multi-context seed constructs. Every syncmer *s*_*i*_ from *S* is used once as the first strobe in the seed construction, with the subsequent strobes chosen using Eq. 2. Both randstrobes and MCS are constructed as described above in the Randstrobes section, with the parameters *w*_min_ and *w*_max_ chosen based on read length.

#### Indexing

The individual MCS are stored as tuples *V*_*i*_ = (*h*_*i*_, *p*_*i*_, *o*_*i*_, *r*_*i*_), where *h*_*i*_ is the MCS hash *h*(*t*_1_, …, *t*_*n*_), *p*_*i*_ is the starting position of the first strobe in the reference sequence, *o*_*i*_ is the offset of the last *n* − 1 strobes (i. e. the distance between the start of the current strobe and the start of the previous strobe), and *r*_*i*_ is the reference index. In memory, *V*_*i*_ takes up 16 bytes: 7 bytes for *h*_*i*_, 4 bytes for *p*_*i*_, 1 byte for *o*_*i*_, and 4 bytes for *r*_*i*_.

The *V*_*i*_ tuples are stored in a vector *V* accompanied by an indexing structure *AEMB index* [32]. We describe it briefly here for self-containment. Given vector *V* of tuples *V*_*i*_ sorted in increasing order by *h*_*i*_ values, and a parameter *b <* 64, the AEMB index defines an array *I*_*b*_, which stores in position *j* the first occurrence position *i* in *V* such that a bit representation of *j* is a *b*-prefix of *i*. If there is no tuple (*h*_*i*_, *p*_*i*_, *o*_*i*_, *r*_*i*_, *f*_*i*_) in *V* such that *j* is a *b*-prefix of *h*_*i*_, a value from the preceding position *I*_*b*_[*j* − 1] is stored instead. We define *I*_*b*_[0] as 0, so a preceding value always exists. Since *V* is sorted by hash values *h*_*i*_, hashes in *V* with the same *b*-prefixes are stored in a single consecutive block. *I*_*b*_ is sometimes referred to as a prefix-lookup vector and has been used in other sequence mapping algorithms [7]. The default value for the parameter *b* depends on the total reference length *Length*, strobe length *s*, and *k*, specifically

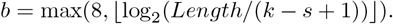

If the hash *h* is queried in *V*, the first position in the block in *V* is the value *I*_*b*_[*h*^*b*^]. The last position of the block is the minimum value in *I*_*b*_ which follows *h*^*b*^ and is different from *I*_*b*_[*h*^*b*^]. The block in *V* can then be searched for the exact match *h* using either linear or binary search.

#### Querying

Our implementation of strobealign with MCS contains all the same algorithm steps as strobealign, with the difference that MCS are used to search for partial matches in case a full match was not found. In more detail, for a query (read), strobealign [37] searches for seed matches in the index, chains the matches (for details, see Suppl. Section 8), and performs an extension alignment step (if strobealign is run in extension-mode). The seeds are constructed from the forward and reverse-complemented read in tuples *V*_*i*_. The rand-strobe version of strobealign differs from the multi-context version in selecting matches from the constructed seeds. In the randstrobe version, the seed is considered a match if its hash is present in the index and if it is not hard masked (i. e. occurs less than 1000 times in the index). In the multi-context version, a seed can be either a full match or a partial match if the full hash value or only the base hash value is present in the index, respectively. The full match criteria are the same as for the randstrobe seed. If a query is not a full match, we search for a base part of a query hash in the index if it was not searched previously. If the base part of a hash is present in the index and is not hard masked, we select it as a partial match. Pseudocode for this procedure can be found in Supplementary Section 1.

Note that since the index *V* is sorted by the hash value, the prefix of the queried hash can also be searched in the index using the same algorithm as for the full hash search, as long as the prefix is longer than the parameter *b*. For MCS, we use this property of the AEMB index to search for *partial matches*, entries in *V* with the same base part of the hash as the query. Thus, looking up the hash of the base (search for partial hit) is cheap as the part of the vector is often in the cache after the full seed has been searched. We refer to the approach described above as strobealign-MCS in the benchmark.

### Bit allocation of the base and the digests

We want to allocate a fraction of bits to represent the base hash value such that it minimizes hash collisions of both full and partial searches. The optimal bit allocation depends on the underlying genome repeat structure, size of *k*, and the number of available bits *B*. However, since the repeat structure is unknown, we aimed to select a reasonable bit partition based on simulated experiments. We computed the fraction of bits allocated to the base hash that minimized the sum of E-hits [37] for full and partial matches with two strobes, for various combinations of *k, w* = |𝒲|, *B*, given a genome size *G* = 2^16^. We investigate both a fully random genome and one with a simulated repeat structure. The details of the experiment are found in Suppl. Section In brief, we found that when *B* is small compared to *G*, we want to allocate more bits to the base. At the same time, for relatively large *B*, the fraction tends towards 0.5 for both genomes (Suppl. Fig. 1). We also observed that the value of *k* in relation to *G* matters since it affects *k*-mer uniqueness. Finally, *w* did not have an impact on the optimal fraction. We acknowledge that biological reference sequences are more repetitive than random strings, and may therefore benefit from using a larger fraction allocated to *B*, compared to that of a random string. We use *B* = 56 in all our benchmarks (and in strobealign), which is sufficient for most genomic databases, and picked a fraction *B* = 40*/*56. The remaining 16 bits are allocated to the digest of the second strobe in 2-order MCS, and 8 bits are allocated to the digests of the second and the third strobe in 3-order MCS.

## Discussion and conclusions

We presented a new seeding approach, multi-context seeds (MCS), that allows multiple layers of matches to be searched. A multi-context seed is a hash value representation of a strobemer, where each strobe in the strobemer is represented by a separate substring of bits in the hash value. This allows the seed to be queried at multiple levels. Particularly, in a sorted array such as the AEMB index (the index used in strobealign), this enables both full and partial matches to be searched in a cache-efficient manner. We show that multi-context seeds produce more matches than *k*-mers and randstrobes with the same subsequence size (Table 1), which indicates increased mapping sensitivity. Additionally, we show that multi-context seeds are as unique as randstrobes and *k*-mers if the full hash is searched (Table 2).

We believe that MCS can be useful in applications where sequence searches at multiple resolutions are useful and where efficiency is needed. We demonstrate the benefit of MCS in one such case by implementing MCS in the read mapper strobealign. We show that strobealign with MCS significantly increases accuracy in single-end and paired-end read alignment and mapping modes compared to strobealign with randstrobes for short read lengths and modest to high variation (SIM4 and SIM6) (Figure 2, Suppl. Fig. S8, S9). Strobealign with MCS has low runtime overhead to strobealign randstrobes, where the overhead depends on the variation levels, while being noticeably faster than BWA-MEM (Figure 3, Suppl. Figs. S10 and S11). BWA-MEM has a slightly faster but much more memory-consuming variant, BWA-MEM2 [43], which we have benchmarked against strobealign [37]. However, optimized implementations for strobealign [45]) also exist that double the speed. MCS is now the default seeding method used in strobealign (v0.17.0). We also implemented a proof-of-concept long-read mapping in strobealign to assess how MCS perform for long-read mapping. We observed that strobealign with MCS was consistently faster than minimap2 while demonstrating comparable and often favorable accuracy, particularly on a human genome (CHM13).

As future work, we aim to use MCS to enable better split-read breakpoint detection with short reads and long-read alignment and to further explore the idea of using MCS for multiple *k*-mer levels. Our experiments of fixing the offset of the second syncmer generated a two-level dynamic *k*-mer index, which showed improvement over other approaches for high diversity datasets (Figure 2, Suppl. Fig. S8 and S9) and to state-of-the-art long-read mapper minimap2 in the proof-of-concept long-read mapping benchmark (Figure 2, Suppl. Fig. S6 and S7). Finally, as our proof-of-concept long-read mapping implementation demonstrated state-of-the-art long-read mapping accuracy, we aim to make strobealign with MCS a complete long-read aligner including piecewise extension alignment and supplementary alignments, features that are necessary for structural variation detection.

## Declarations

### Availability of data and materials

Strobealign 0.17.0 is available under the MIT license from https://doi.org/10.5281/zenodo.17974973 and the development version can be found at https://github.com/ksahlin/strobealign. The bit allocation analysis is found at https://github.com/ksahlin/misc/tree/main/mcs. The code for the sequence matching results is located at https://github.com/Itolstoganov/strobemers/tree/multi-context-benchmarks. The code for the uniqueness benchmark is available at https://github.com/Itolstoganov/strobealign/tree/uniqueness_benchmark. The code for the read alignment experiments is available at https://github.com/NBISweden/strobealign-evaluation/. Biological datasets BIO150 (Illumina WGS 2×150bp, HG004) and BIO250 (Illumina WGS 2×250bp, HG004) analyzed in this study are found at https://github.com/genome-in-a-bottle/giab_data_indexes.

## Acknowledgements

The computations were enabled by resources provided by the National Academic Infrastructure for Supercomputing in Sweden (NAISS), partially funded by the Swedish Research Council through grant agreement no. 2022-06725.

## Authors’ contributions

KS and IT conceived the concept of multi-context seeds. IT, MM, and KS implemented MCS in strobealign. IT analyzed the seed constructs using uniqueness and coverage metrics. MM and IT benchmarked strobealign-MCS against state-of-the-art alignment tools and analyzed the results. NB implemented chaining in strobealign and wrote the supplementary section on chaining. IT and KS performed the variant calling analysis. IT and KS drafted the paper with input from MM. All authors read and contributed to the final manuscript.

## Ethics approval and consent to participate

Not applicable.

## Consent for publication

Not applicable.

