## Supplementary file for "Multi-context seeds enable fast and high-accuracy read mapping"

### 1 Pseudo code for querying MCS in strobealign

**Algorithm 1:** Querying MCS

**input :** Query MCS ($Q$), Index ($R$), Total bits for hash value ($B$), bit allocation for base hash ($b$)
**output:** Matches

```
Initialize vector of matches M
Initialize set of previously searched partial seed hash values P
Function findMatches(Q, R)
    for q ∈ Q do
        m_full = R.FindBitMatches(q[0 : B], R)
        if ∃ m_full then
            M.push_back(m_full)
        else
            if q[0 : B − b] ∉ P then
                m_partial = R.FindBitMatches(q[0 : B − b], R) //Same location in R as full seed
                if ∃ m_partial then
                    M.push_back(m_partial)
                    P.add(q[0 : B − b])
    return M
```

### 2 Bit allocation analysis

The uniform genome was simulated by sampling nucleotides uniformly at random. Therefore, it contains no repeat structure beyond what is expected for small $k$. For this genome, most $k$-mers of size 8 ($4^8 = 2^{16}$ $k$-mers) are expected to be occurring once in a uniform genome of size $2^{16}$. For the repetitive genome, we simulated an origin fragment of length 512 nucleotides. We then simulated mutated copies (substitutions) of this fragment (5% mutation rate) and appended it to the genome. The repeated genome therefore contains 128 mutated repeat copies of the fragment where the largest $k$-mer size of 8 cannot be expected to be largely unique.

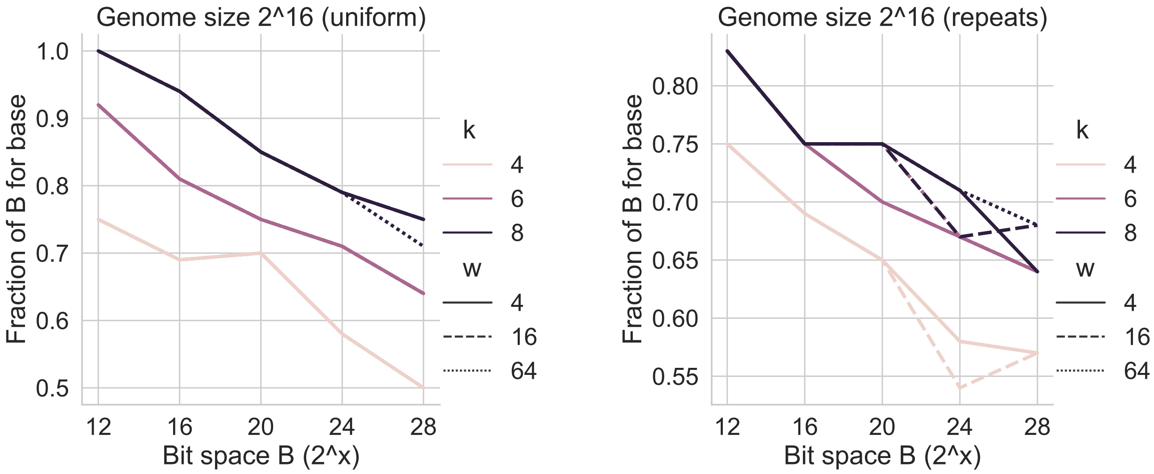


Fig. S1: Fraction of bits (y-axis) that minimizes E-hits for the sum of full and partial hits, for various sizes of bit allocations $B$ for the seed (x-axis).

### 3 Metrics for the sequence matching experiment

For all the evaluated seed constructs, we say that a *match* between two sequences $s$ and $s'$ occurs at positions $i$ of $s$ and $i'$ of $s'$ if the seed starting in position $i$ has the same hash value as the seed of $s'$ starting in position $i'$. Additionally, if two MCS have the same hash value prefixes corresponding to their first $m$ strobes, we say that a *partial match* of rank $m$ occurs at positions $i$ of $s$ and $i'$ of $s'$. We refer to the fraction of match starting positions as the *match percentage*. In the case of MCS which can have both full and partial matches, the match percentage is the non-redundant sum of full and partial matches. That is, if a position in $s$ is counted as a full match for the purpose of match percentage, it cannot be counted as a partial match. Similarly, if a position is counted as a partial match of rank $m$, it cannot be counted as a partial match of a smaller rank.

For a $k$-mer, we say that a match occurring at position $i$ produces a *coverage subsequence* over positions $[i, i + k - 1]$. A randstrobe match occurring at position $i_1$ produces a coverage over a subsequence of $s$ corresponding to the strobes that constitute the matching randstrobe. Specifically, if a randstrobe consists of $n$ strobes starting at positions $i_1, i_2, \ldots, i_n$, its coverage subsequence over $s$ is $[i_1, i_1 + s - 1] \cup [i_2, i_2 + s - 1] \cup \cdots \cup [i_n, i_n + s - 1]$. A multi-context match produces the same coverage subsequence if it is a full match, and the coverage subsequence over the first $m$ strobes if it is a partial match of rank $m$. Given the coverage subsequence $s_{i_1}, s_{i_2}, \ldots, s_{i_m}$ of a match, we say that the *match coverage* of this match constitutes the interval $[i_1, i_m]$, and the *sequence coverage* constitutes the set of positions $i_1, i_2, \ldots, i_m$. The total sequence coverage of $s$ is defined as the percentage of $s$ length covered by the union of the sequence coverage positions of all matches and the total match coverage of $s$ as the percentage of the string covered by the union of the match coverage positions.

Finally, in addition to the match percentage, total sequence coverage, and total match coverage, we employ the island E-size metric. As in [1], we define an *island* as a continuous interval of positions not covered by matches. If $X$ is a set of islands in a string $s$, and $|x|$ is the length of an island interval, the *island E-size* $E$ is defined as $E = \frac{1}{|s|} \sum_{x \in X} |x|^2$.

| | | SIM-R | | | | | | | | | | | |
|---|---|---|---|---|---|---|---|---|---|---|---|---|---|
| | | 0.01 | | | | 0.05 | | | | 0.1 | | | |
| | | m | mc | sc | E-size | m | mc | sc | E-size | m | mc | sc | E-size |
| kmers | 30 | 74.5 | 96.0 | 96.0 | 1.1 | 22.3 | 54.6 | 54.6 | 43.7 | 4.7 | 18.2 | 18.2 | 289.8 |
| kmers | 10 | 90.4 | 99.2 | 99.2 | 0.0 | 62.3 | 91.2 | 91.2 | 0.9 | 38.8 | 75.8 | 75.8 | 4.2 |
| kmers | 15 | 86.7 | 98.6 | 98.6 | 0.1 | 48.3 | 81.7 | 81.7 | 4.2 | 22.6 | 54.0 | 54.0 | 22.3 |
| kmers | 20 | 82.4 | 97.9 | 97.9 | 0.3 | 37.3 | 72.6 | 72.6 | 11.0 | 13.4 | 38.6 | 38.6 | 62.6 |
| randstrobes | (2, 15, 25, 50) | 70.7 | 99.9 | 98.2 | 0.0 | 18.1 | 87.8 | 72.7 | 8.2 | 3.4 | 44.7 | 31.2 | 117.0 |
| randstrobes | (3, 10, 25, 50) | 66.7 | 100.0 | 98.8 | 0.0 | 14.7 | 98.2 | 78.3 | 1.1 | 2.5 | 67.0 | 33.8 | 68.5 |
| multi-context | (2, 15, 25, 50) | 86.7 | 99.9 | 98.6 | 0.0 | 48.3 | 91.8 | 81.6 | 2.8 | 22.6 | 62.6 | 53.9 | 20.9 |
| multi-context | (3, 10, 25, 50) | 91.2 | 100.0 | 99.4 | 0.0 | 62.9 | 99.7 | 92.2 | 0.1 | 39.3 | 93.9 | 77.0 | 1.6 |

Table S1: Match statistics for $k$-mers, randstrobes, and MCS under mutation rates of 0.01, 0.05, and 0.1. Here, $m$ denotes the number of matches (both full and partial for MCS) as a percentage of the total number of extracted subsequences for the protocol, *sc* (sequence coverage) and *mc* (match coverage) are shown as the percentage of the total sequence length, and *E-size* is the expected island size. The randstrobes and multi-context seeds were constructed with the window parameters $w_{\min} = 25$ and $w_{\max} = 50$

### 4 Parameters used in strobealign

Parameters $k$, $s$ (subsampling of syncmers), and $w_{\min}$, $w_{\max}$ (window for selection of second strobe) typically need to be chosen depending on the input read length. When run, strobealign therefore estimates the read length $\widetilde{x}$ from the first 500 reads and picks one of the following parameter combinations depending on $\widetilde{x}$. These parameters were chosen with the help of a grid search optimization for common read lengths (https://github.com/ksahlin/strobealign/pull/345).

$$(k, s, w_{\min}, w_{\max}) = \begin{cases} (18,14,1,4), & \text{if } \widetilde{x} \leq 70 \\ (20,16,1,6), & \text{if } 70 < \widetilde{x} \leq 90 \\ (20,16,2,6), & \text{if } 90 < \widetilde{x} \leq 110 \\ (20,16,3,8), & \text{if } 110 < \widetilde{x} \leq 135 \\ (20,16,5,11), & \text{if } 135 < \widetilde{x} \leq 175 \\ (22,18,6,16), & \text{if } 175 < \widetilde{x} \leq 375 \\ (23,17,5,15), & \text{if } 375 < \widetilde{x}. \end{cases} \quad (1)$$

In strobealign's command-line interface and documentation, $w_{\min}$ and $w_{\max}$ are called `l` and `u`, respectively (lower and upper).

In the *noisy* profile, parameters $(k, s, w_{\min}, w_{\max}) = (16, 12, 2, 2)$ are used independent of read length.

### 5 Read simulations

#### 5.1 Short reads

The simulated SIM0, SIM1, SIM4 and SIM6 datasets were generated using the Snakemake [5] workflow at https://github.com/NBISweden/strobealign-evaluation. Each dataset consists of reads from four genomes (fruit fly, maize, CHM13 and rye). For each dataset and genome, the following libraries were generated: 1 million paired-end reads of read lengths 75, 100, 150, 200, 300 and 500 nt, as well as single-end reads of lengths 1000, 5000 and 10 000 nt (with read counts 500 000, 100 000 and 50 000 respectively).

Read simulation is a two-step process: First, variants are generated (single-nucleotide variants and small indels, using `mason_variator`), and then reads with sequencing errors are simulated using `mason_simulator`. We use four categories of datasets; SIM0, SIM1, SIM4, and SIM6. An increasing number indicates higher variation. They represent:

**SIM0:** No variation. No read errors.
**SIM1:** No variation. Illumina read error rates.
**SIM4:** Variation `--snp-rate 0.005 --small-indel-rate 0.0005 --max-small-indel-size 50`. Illumina read error rates.
**SIM6:** Variation `--snp-rate 0.05 --small-indel-rate 0.002 --max-small-indel-size 100`. Illumina read error rates.

Thus, SIM0 reads are substrings of the reference. They were generated with the `readsimulator.py` script found in the repository. SIM0 serves as a baseline and sanity check (all reads should be mappable, and all uniquely mapping reads should be mapped to the correct location).

SIM1, SIM4 and SIM6 consist of simulated Illumina reads that were generated with `mason_simulator`. SIM1 was only used for assessing whether Illumina-like error rates served reasonable proxy to HiFi simulated reads in our long-read mapping benchmark (see Suppl. Section 5.2). SIM4 and SIM6 contains variations simulated with `mason_variator` with parameters given above.

For read lengths 300 nt and 500 nt, `mason_simulator` was run with `--fragment-mean-size 700` (the default fragment size is otherwise 300). For read length 1000 nt and longer, `mason_simulator` was run with `--fragment-mean-size` set to 1.5 times the read length. (This is required even though only single-end reads were simulated.)

Single-end reads were obtained by leaving out the R2 reads from the simulated paired-end libraries.

A crash in `mason_simulator` when simulating reads for the large rye genome was fixed after we reported it. To use the fixed version, we compiled the software from the SeqAn repository (https://github.com/seqan/seqan, tag `seqan-v2.5.0rc2`).

Missing numbers (SIM2, SIM3, and SIM5) indicate variation levels that have previously been used in [7] and in an earlier version of this study [8]. We believe SIM0, SIM4, and SIM6 represent a broader span of divergence levels to previous levels.

#### 5.2 Long reads

We also used Mason to simulate long reads. We used Mason as it can simulate longer reads with ground truth from genomes with variation (according to SIM mutation rates described above), unlike some other long read simulators we tried (e. g., PBSIM [6]).

The other tools we tried can only simulate reads from a reference without variation, so we opted to use mason simulator to simulate long reads with an Illumina error profile (SIM1) as a proxy for HiFi reads, in order to assess long read mapping on genomes with different variation levels. However, we first also performed a test as to whether results with Mason-simulated long reads mimicked the performance of long-read mapping using HiFi simulated reads with PBSIM on a genome without variation. We confirm similar results between the two read sets (Suppl. Figs. S2 and S3). Hence, in this proof-of-concept long-read mapping evaluation we believe mason simulations serves a good proxy to HiFi read error rates. The PBSIM benchmark is described in the following paragraph.

**PBSIM HiFi reads evaluation** We simulated PacBio circular consensus (HiFi) reads using the Snakemake workflow [5] available at `https://github.com/NBISweden/strobealign-evaluation`. We generated **HIFI** read datasets with read lengths of 1 kbp, 5 kbp, and 10 kbp, consisting of 500,000, 100,000, and 50,000 reads, respectively. First, PacBio continuous long reads (CLR) were simulated using `pbsim3` [6] with the following parameters: `--strategy wgs --method qshmm --qshmm QSHMM-RSII.model --sd-length 0`. Circular consensus reads were then generated from the simulated CLR reads using `ccs` version 6.4.0 (`https://github.com/PacificBiosciences/ccs`) with default settings.

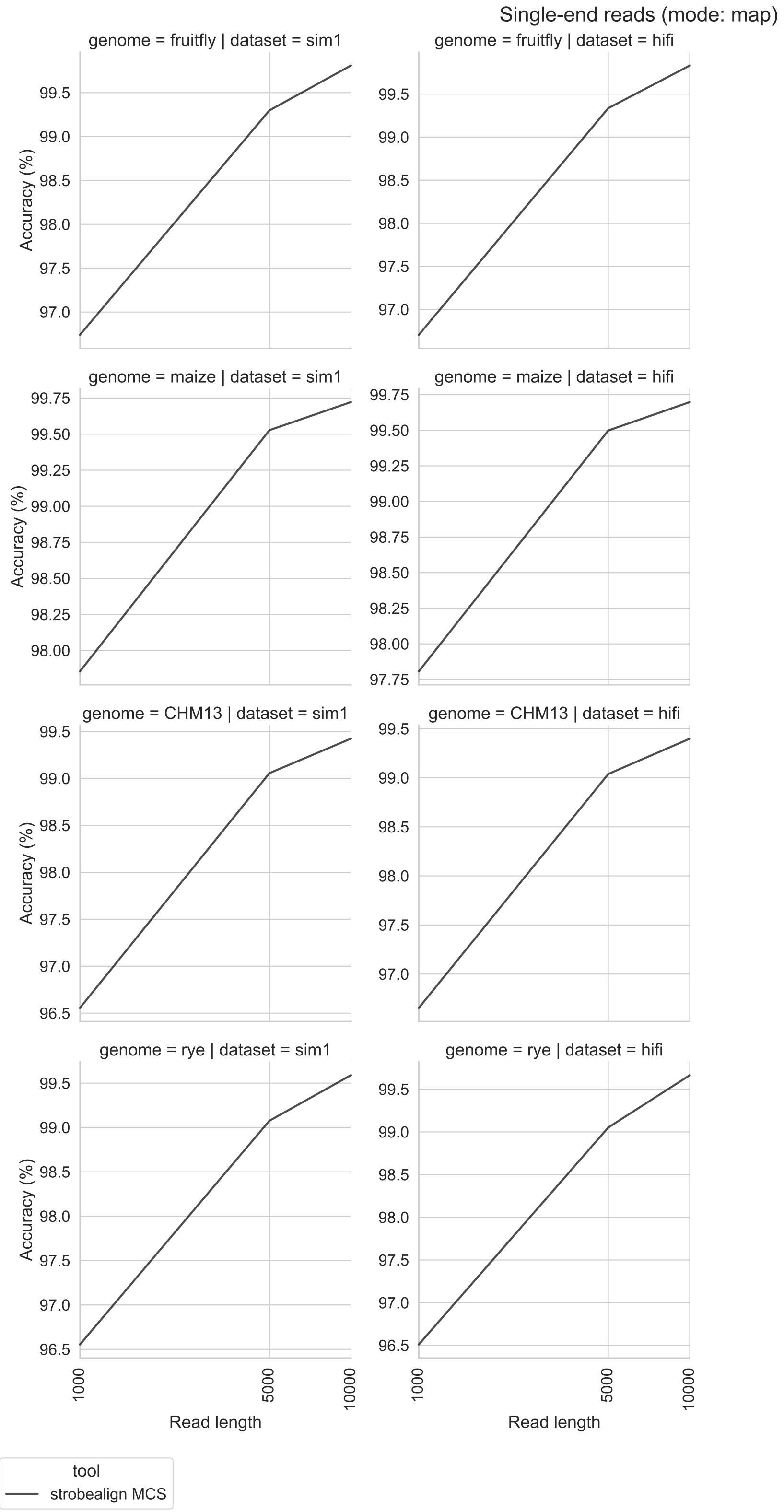


Fig. S2: Accuracy of single-end reads simulated from the fruit fly (top row), maize (second row), CHM13 (third row), and rye (bottom row) for the **SIM1** and **HIFI** datasets.

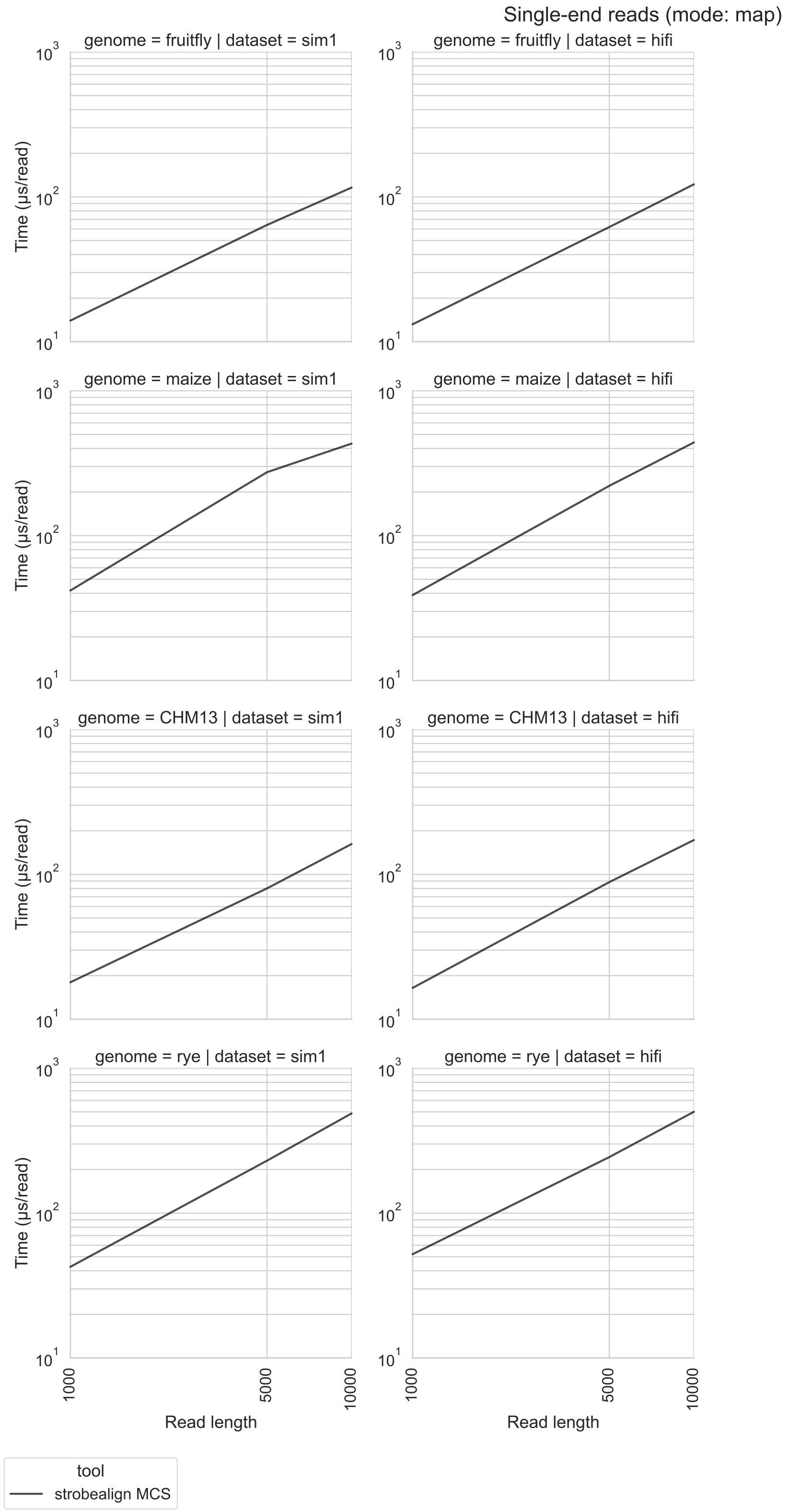


Fig. S3: Runtime of the read mapping for the **SIM1** and **HIFI** datasets. simulated from the fruit fly (top row), maize (second row), CHM13 (third row), and rye (bottom row).

### 6 Read alignment benchmark for the rye, fruit fly, and maize genomes

The read alignment benchmark for this study can be reproduced using the workflow at `https://github.com/NBISweden/strobealign-evaluation` in the `mcs/` folder. Tables with our measurements are available in CSV format from the same location.

The benchmarks were run on a PC with an AMD Ryzen 7 PRO 5850U CPU with 8 cores. Each program was allowed to use 8 threads.

Minimap2 2.30, BWA-MEM 0.7.19 and X-Mapper 1.2.0 were installed from Bioconda [3], and strobealign 0.17.0 was compiled from sources.

Strobealign and minimap2 were run in both alignment mode (producing SAM output) and mapping-only mode (producing PAF output). BWA-MEM and X-Mapper do not have a mapping-only mode.

The runtime shown for all tools excludes indexing. A pregenerated index (stored on disk) was used for BWA-MEM and minimap2.

#### 6.1 X-Mapper evaluation

We made an effort to compare strobealign with the recently published [2] X-Mapper tool, version 1.2.0. The program has a few limitations that we had to work around before we could plug it into our evaluation pipeline. We list below the limitations and our workarounds and assumptions.

1. The main one is that X-Mapper often outputs multiple alignments for a read without marking one of them as the primary one. The SAM specification requires one alignment to be primary and additional alignments to be marked as secondary. To make the output comparable with that of the other tools, we first sort the alignments for a read (or read pair) by score (according to the `AS` tag) and, if there is a tie, randomly choose one as the primary one. This is the same strategy other read mappers including strobealign employ when they find multiple equally good alignments. Our accuracy computation script skips secondary alignments.
2. X-Mapper does not output aligned reads in the same order as the input reads. Since this is required by our evaluation script, we added an extra step that sorts reads by name (`samtools sort -n`). We do not include this step in the runtime measurement.
3. Alignments for some reads are missing from the output. We assume these are unmapped reads and treat them accordingly in our evaluation script.
4. Since X-Mapper appears to not automatically infer insert size when mapping paired-end reads, we provided an appropriate value using the `--spacing` option based on the target fragment size used during read simulation.

We list further noteworthy behaviors of this version of X-Mapper that were irrelevant for the benchmarks but could be an issue in a production setting:

- Reference contigs in the SAM header are in a different order than in the input FASTA file.
- For FASTQ input, the base qualities are omitted in the output SAM and replaced with an asterisk `*`.
- The mapping quality is always set to either 0 or 255.
- The alignment score (`AS`) SAM tag has the type "f" (floating point number) instead of "i" (integer) as required by the SAM specification.

We do not show memory usage in the plots because the maximum resident set size (RSS) that we measure would give the wrong impression for this program as it runs on the Java Virtual Machine (JVM). When provided with a maximum amount of memory to use, the JVM tends to fill it even when a smaller amount would work just as well. All runs of X-Mapper finished successfully with option `-Xmx48g` (48 GB maximum memory usage), which therefore serves as an upper bound for the minimum memory usage.

Since X-Mapper indexes the reference on-demand while mapping, separating indexing and mapping runtime is hard. As a conservative estimate, we consider the time it takes until at least 10 000 reads have been mapped to be the indexing time and the rest the mapping time.

X-Mapper cannot persist its index and instead recomputes it on every invocation. Indexing takes about 15 minutes for CHM13 using 8 threads. For comparison, strobealign needs about 35 seconds.

Figure S4 shows how X-Mapper performs in terms of accuracy and runtime on the CHM13 genome in comparison to strobealign-MCS.

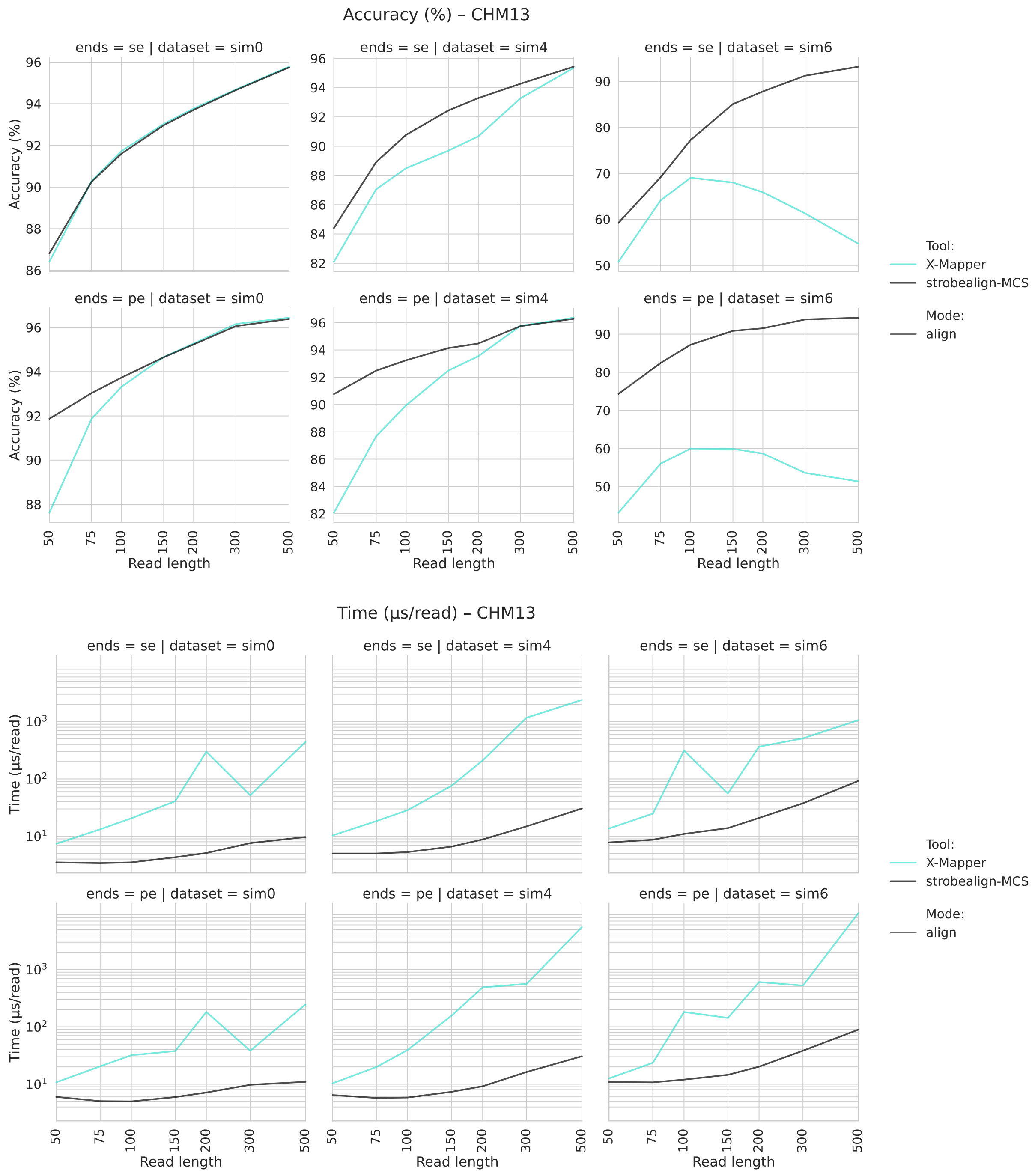


Fig. S4: X-Mapper accuracy and runtime on CHM13 compared to strobealign-MCS (single-end and paired-end reads).

#### 6.2 Seed lookup strategies

The index data structure introduced for multi-context seeds allows full and partial hash lookups. This enables us to implement different seed lookup strategies: randstrobes, MCS, and $k$-mer only. The default, as discussed, is strobealign-MCS, which first looks up the full hash and, if the lookup fails, falls back to a partial hash lookup. Strobealign randstrobes always performs a full lookup only.

Strobealign with $k$-mer-only always performs a partial lookup, ignoring the full hash. This allows us to simulate purely $k$-mer-based seeds (i. e., syncmers in strobealign).

In strobealign 0.17.0, the seed lookup strategy can be chosen on the command-line as follows: `--mcs=always` for strobealign-MCS (the default), `--mcs=first-strobe` for strobealign $k$-mers, and `--mcs=off` for strobealign randstrobes.

Figure S5 compares the performance of the three options.

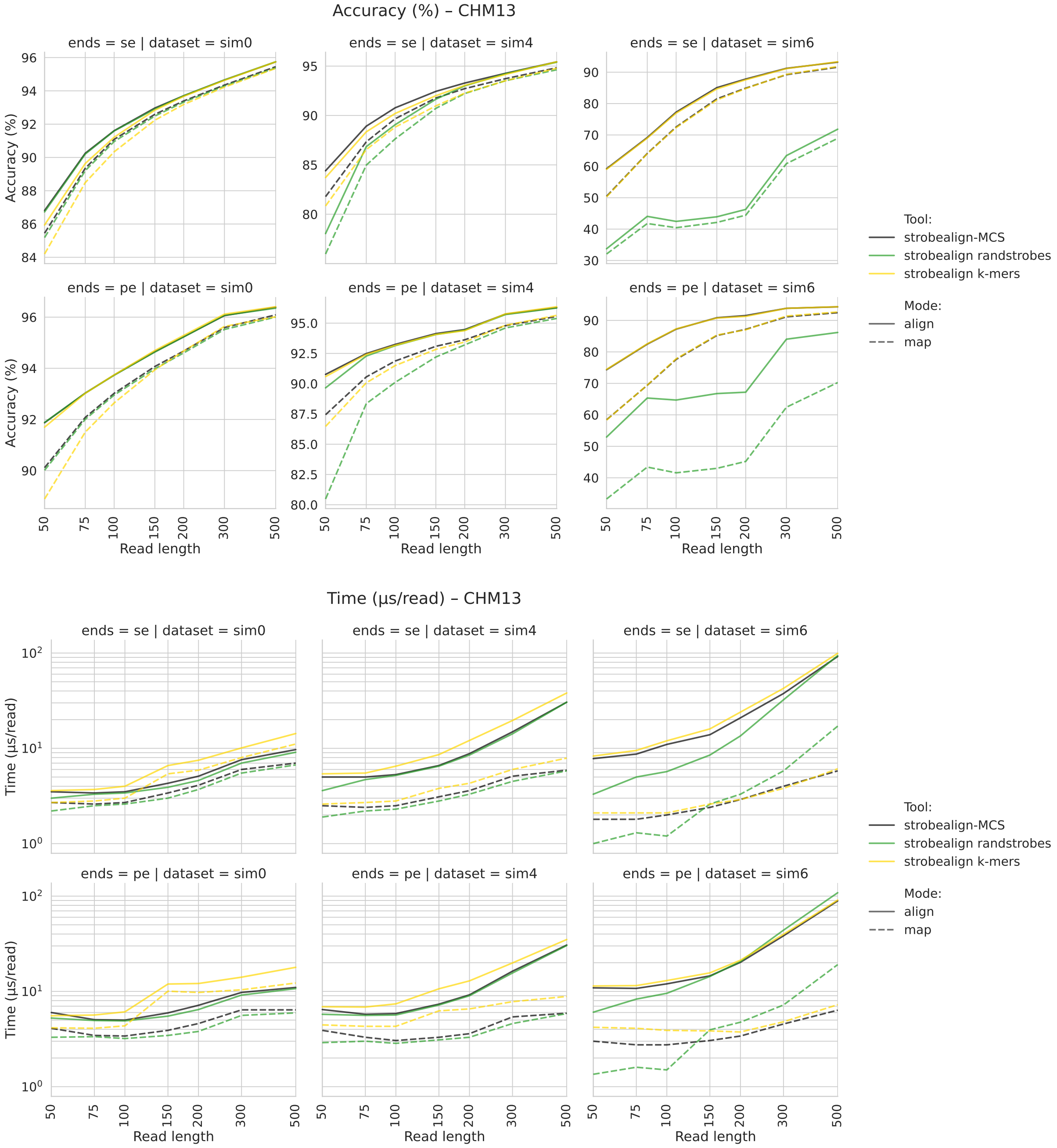


Fig. S5: Accuracy and runtime for the three seed lookup strategies in strobealign: MCS, randstrobes and $k$-mer only. Results for the other genomes are comparable (not shown).

#### 6.3 Benchmark results for long reads

Note these are mapping-only results (PAF output). Support for base-level alignment (SAM output) for long reads is under development in strobealign.

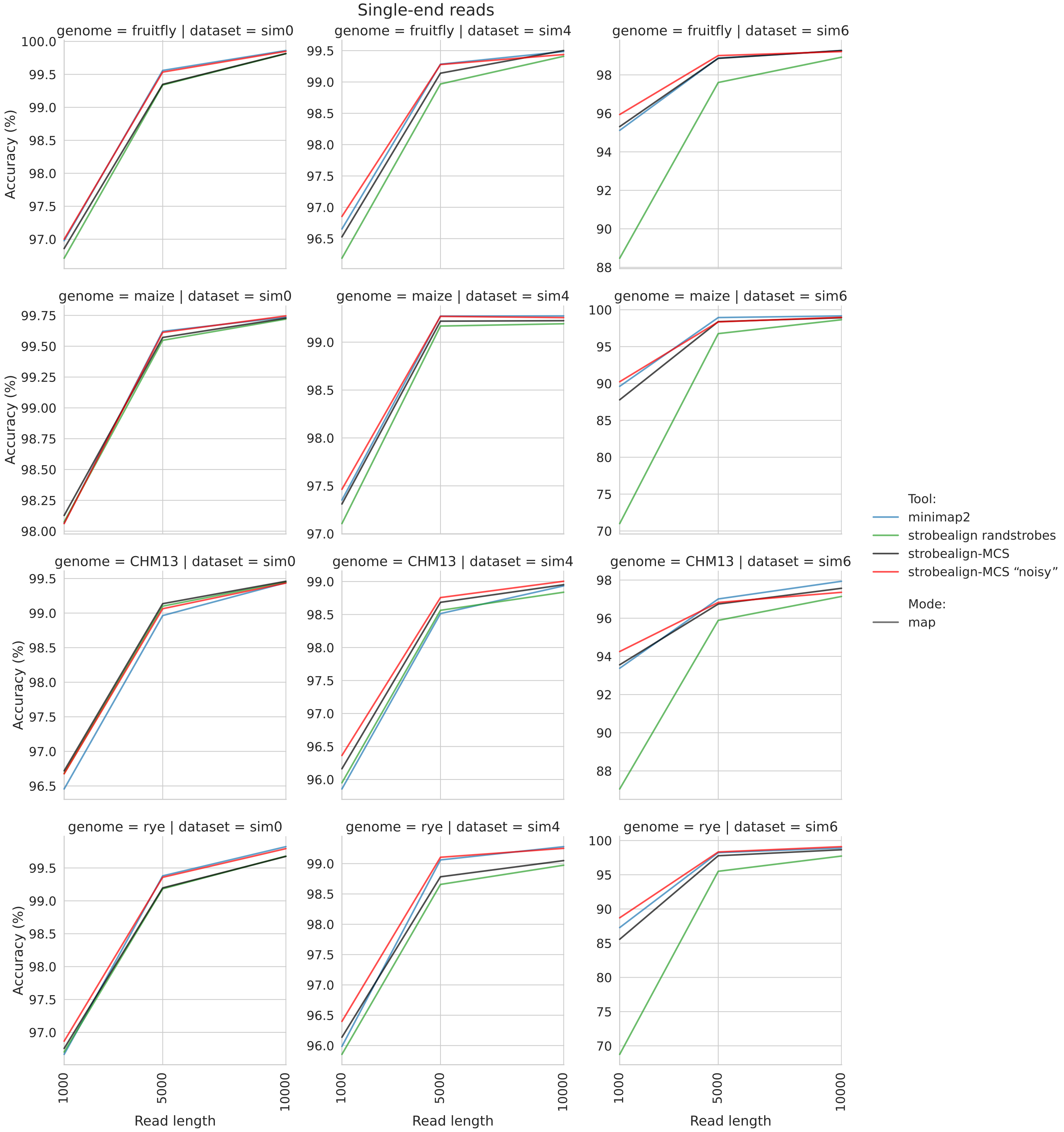


Fig. S6: Accuracy of strobealign on long reads

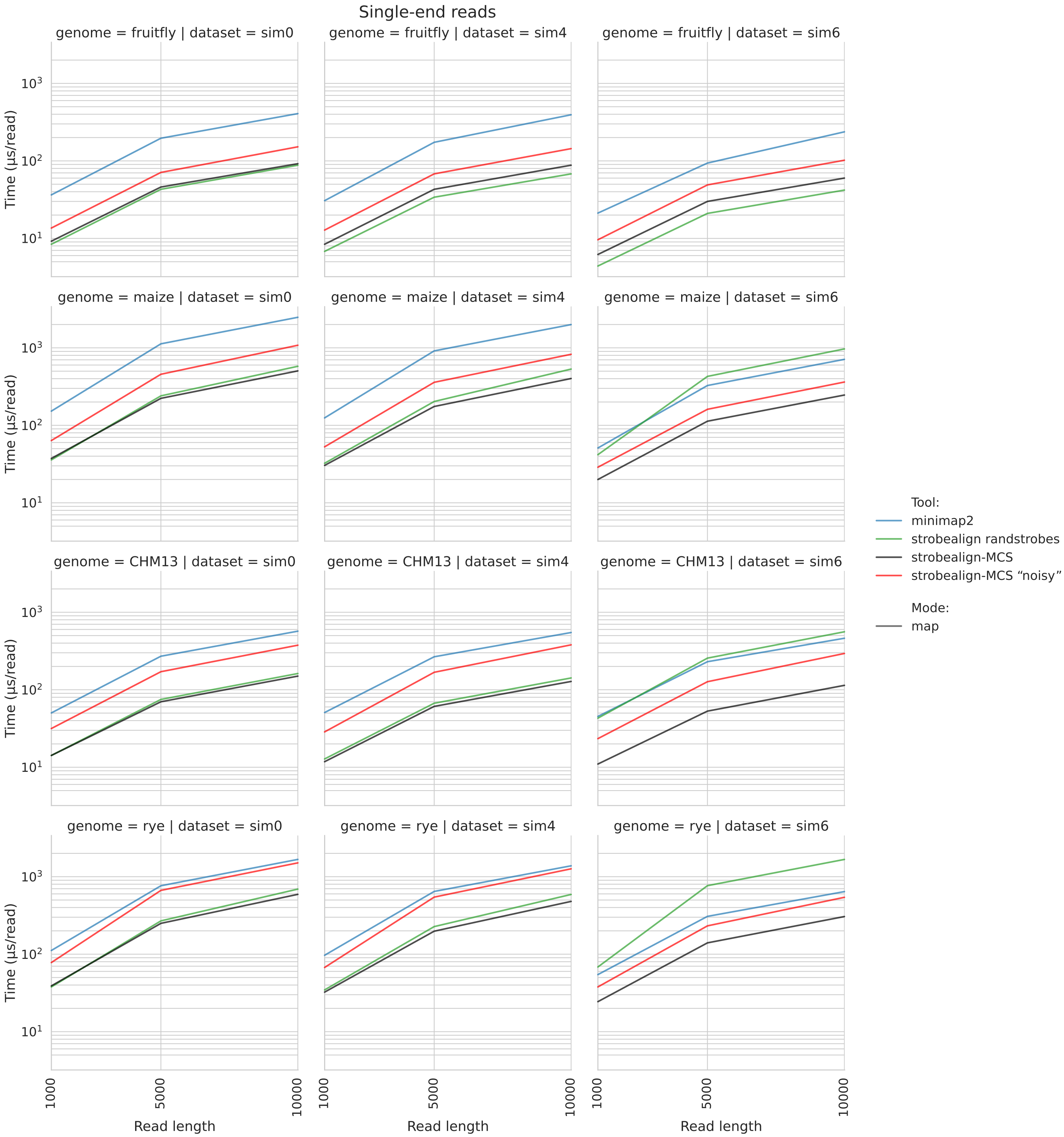


Fig. S7: Runtime of strobealign on long reads

#### 6.4 Full benchmark results (all four genomes) for short reads

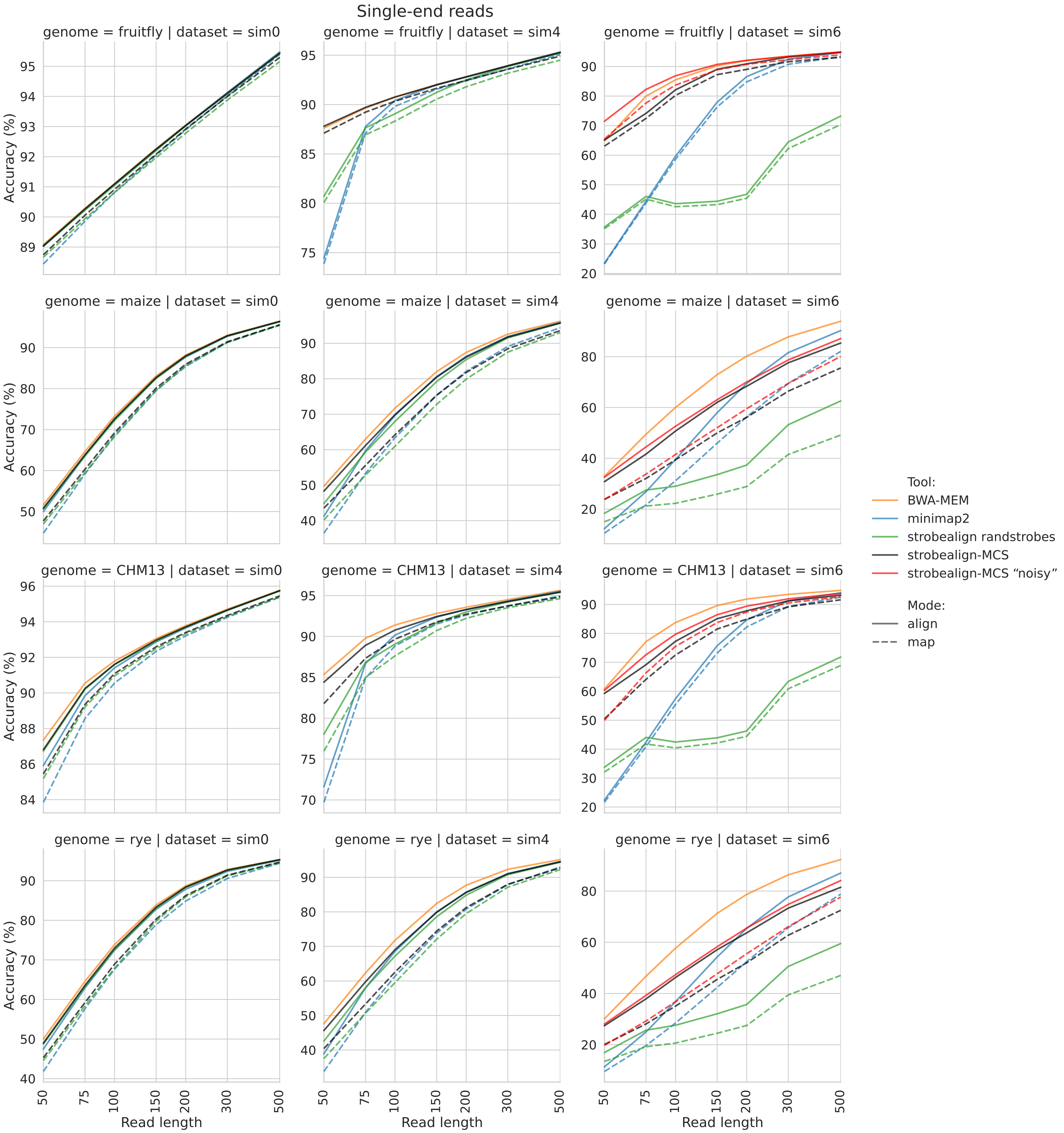


Fig. S8: Accuracy of single-end reads simulated from the fruit fly (top row), maize (second row), CHM13 (third row), and rye (bottom row) genomes for the **SIM0** (left), **SIM4** (middle), and **SIM6** (right) datasets

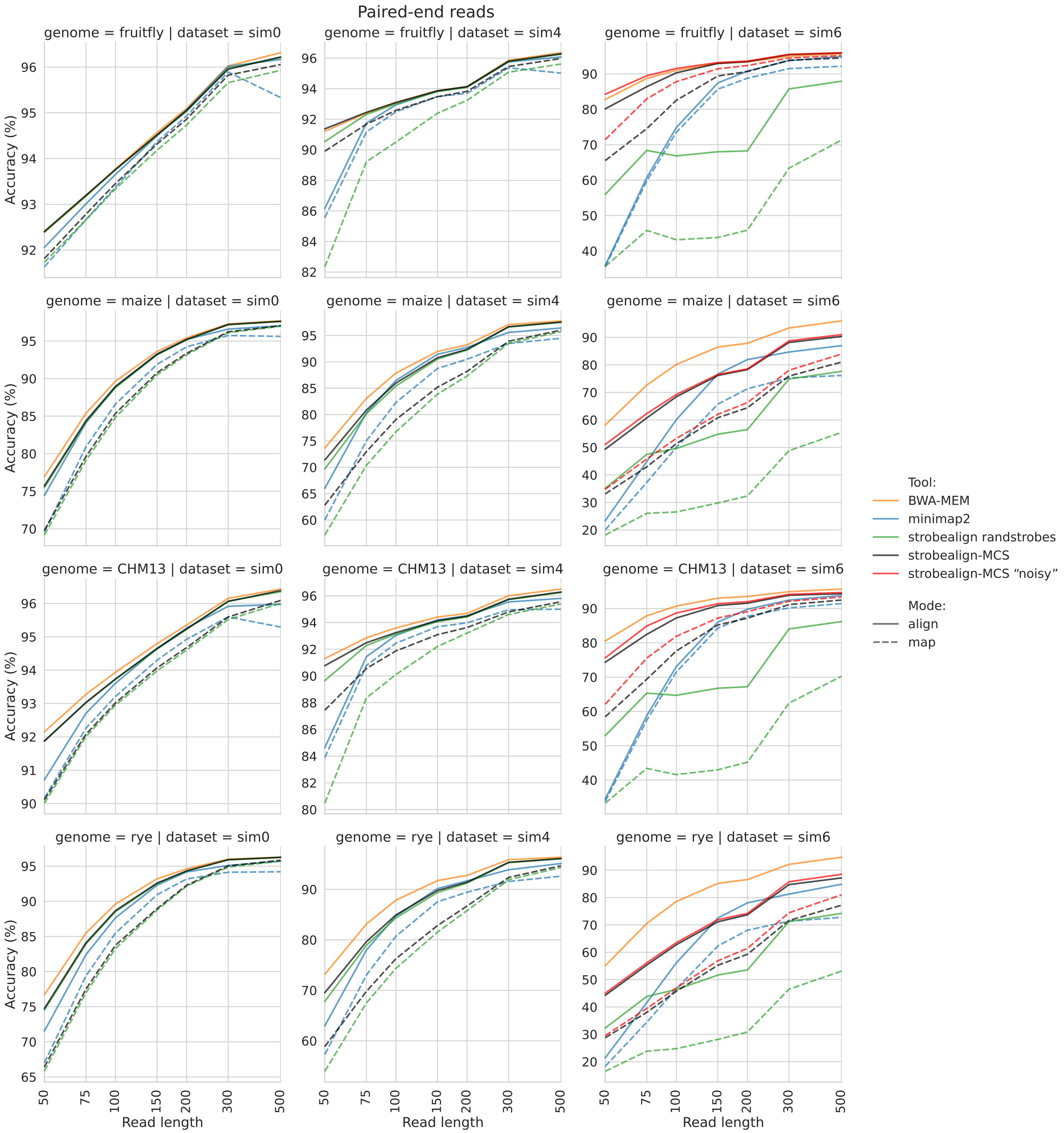


Fig. S9: Accuracy of paired-end reads simulated from the fruit fly (top row), maize (second row), CHM13 (third row), and rye (bottom row) genomes for the **SIM0** (left), **SIM4** (middle), and **SIM6** (right) datasets

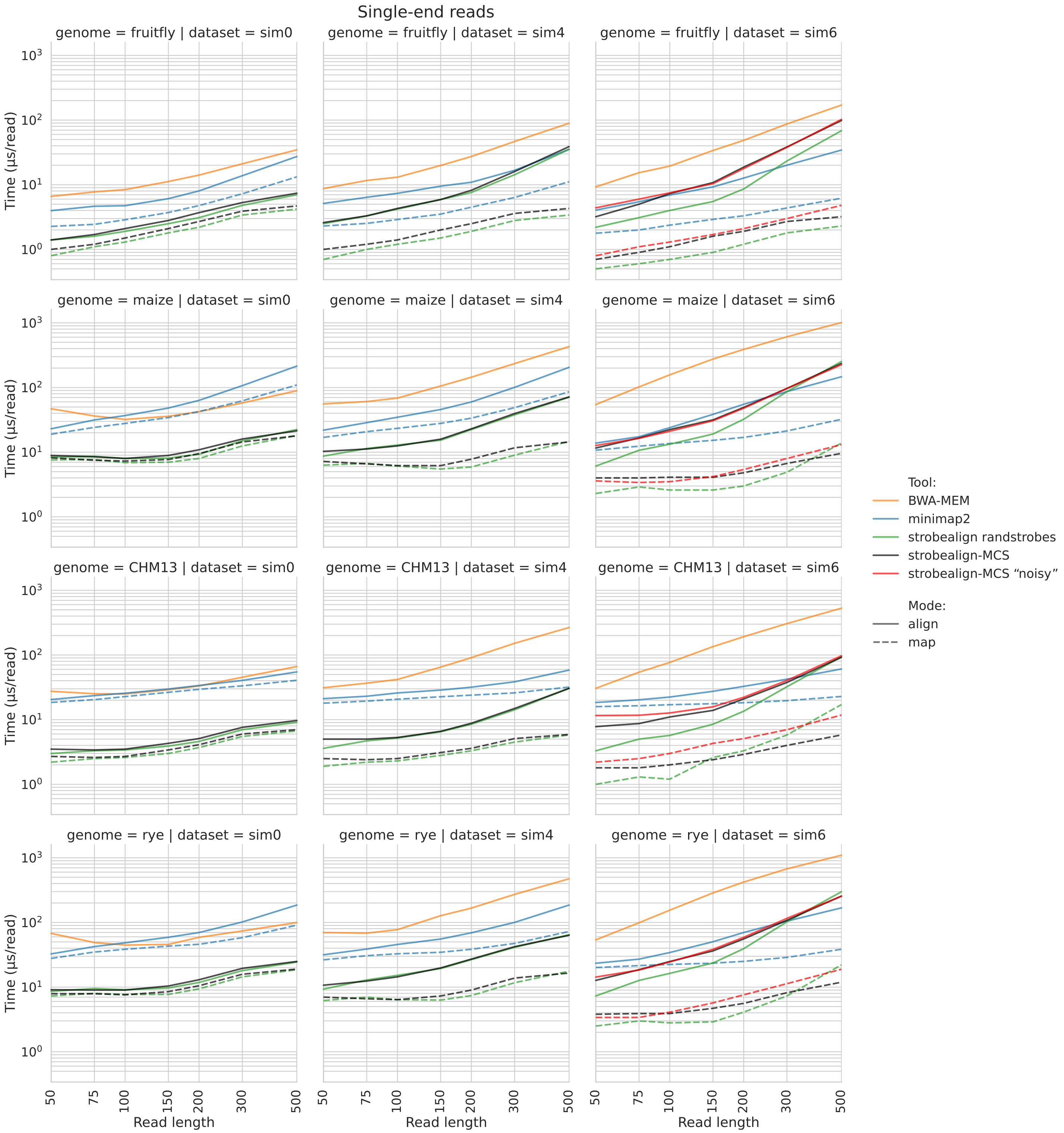


Fig. S10: Runtime of the single-end read alignment for the **SIM0** (left), **SIM4** (middle), and **SIM6** (right) datasets. Reads were simulated from the fruit fly (top row), maize (second row), CHM13 (third row), and rye (bottom row) genomes.

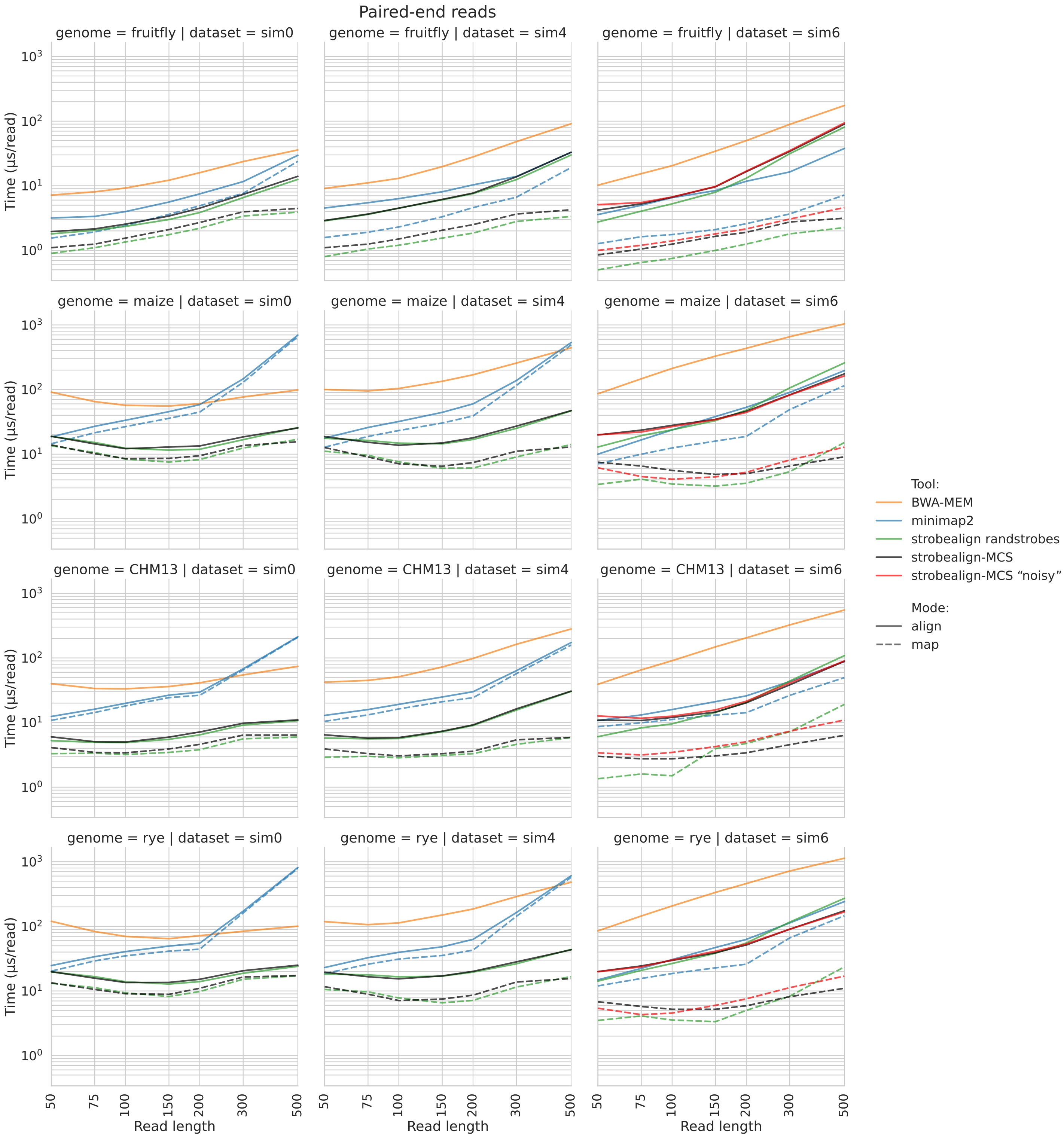


Fig. S11: Runtime of the paired-end read alignment for the **SIM0** (left), **SIM4** (middle), and **SIM6** (right) datasets. Reads were simulated from the fruit fly (top row), maize (second row), CHM13 (third row), and rye (bottom row) genomes.

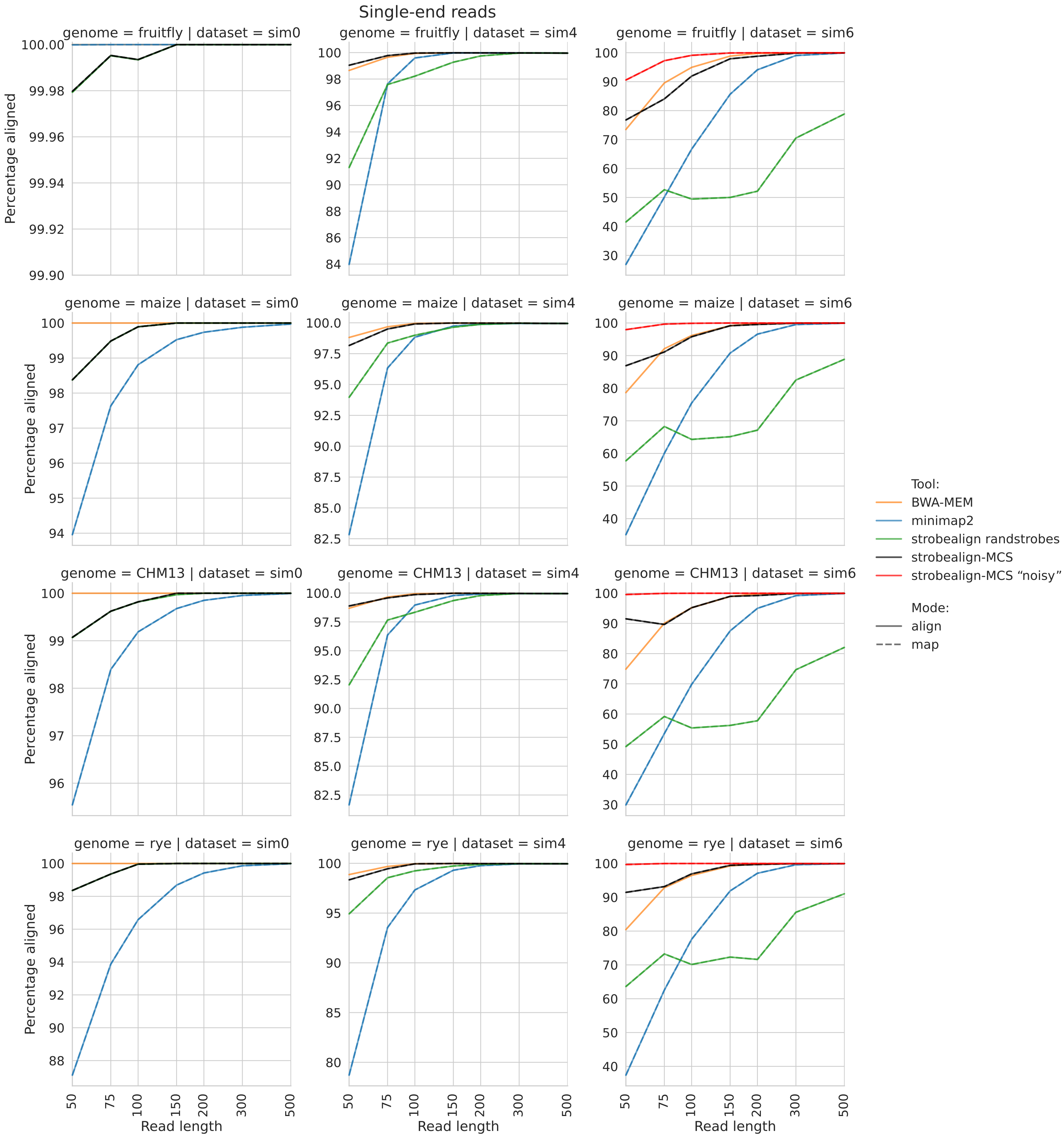


Fig. S12: Mapping rate of single-end reads simulated from the fruit fly (top row), maize (second row), CHM13 (third row), and rye (bottom row) genomes for the **SIM0** (left), **SIM4** (middle), and **SIM6** (right) datasets

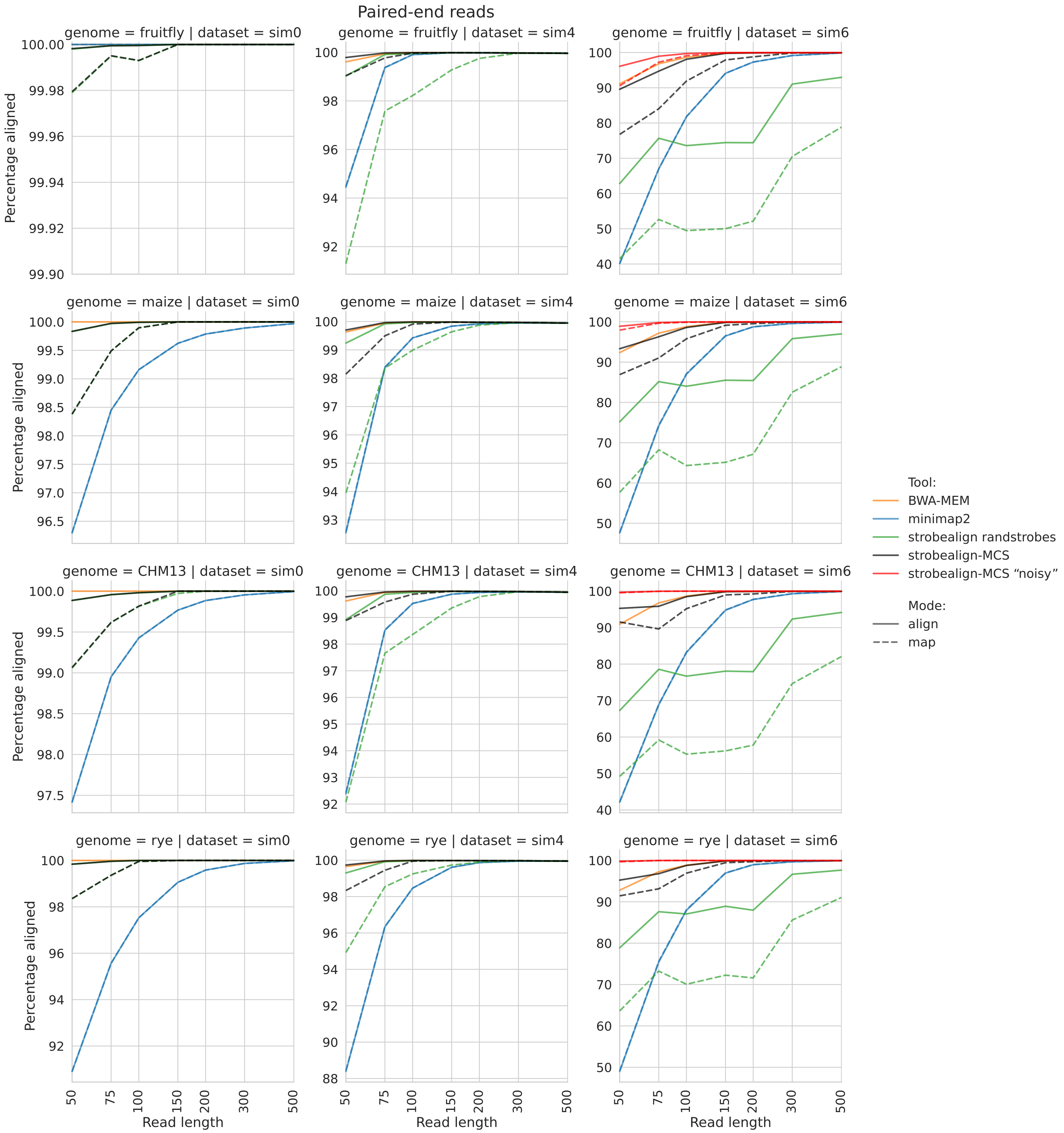


Fig. S13: Mapping rate of paired-end reads simulated from the fruit fly (top row), maize (second row), CHM13 (third row), and rye (bottom row) genomes for the **SIM0** (left), **SIM4** (middle), and **SIM6** (right) datasets

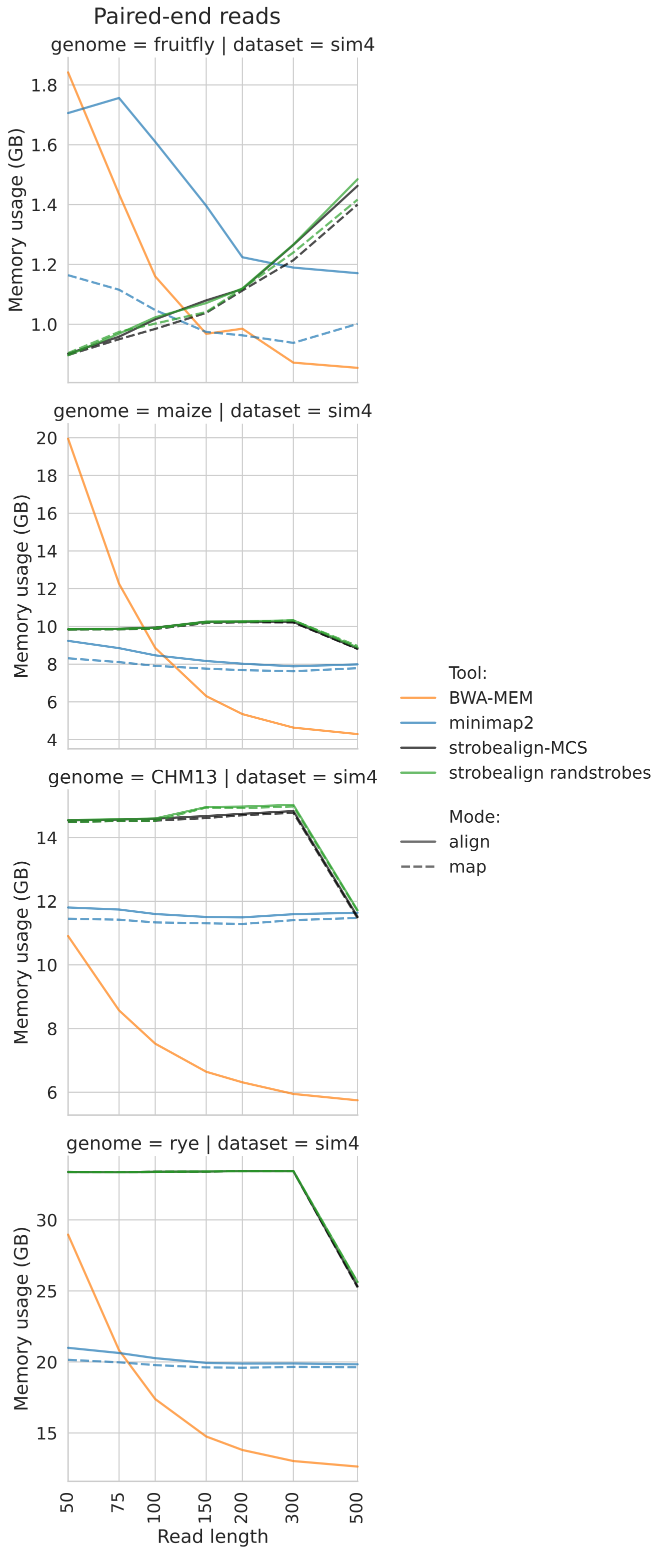


Fig. S14: Memory usage. Only paired-end memory usage on SIM4 is shown as memory usage for single-end reads and for the other datasets is nearly identical.

### 7 3-strobe evaluation

While the current version of strobealign utilizes multi-context seeds consisting of two strobes, we implemented an additional version which employs three-strobes multi-context seeds, with an additional layer of search. This version can be accessed at `https://github.com/ksahlin/strobealign/tree/3-strobes-paper`.

We evaluated this version in three modes with different querying procedure. The (3|2|1) mode searches for the full seed (which represents all three strobes) in the index. If the full seed was not found, the partial seed consisting of the first two strobes are searched, followed by the first strobe search if the search for the partial seed was not successful. The (3|2) and (3|1) modes similarly start by searching the full seed, followed by the search for the first two strobes or the first strobe, respectively.

We evaluated the 3-strobe version on the fruitfly, maize, and CHM13 datasets in three querying modes, along with the 2-strobe version. The evaluation was performed in mapping-only mode. The $k$, $s$, $w_{\min}$, and $w_{\max}$ parameters are the same as in the 2-strobe version. Out of the 56-bit hash, 40 bits were allocated for the first strobe, and 8 bits for the second and third strobe, respectively.

As shown in Figure S15, (3|2|1), (3|2), and 2-strobe modes demonstrate similar accuracy across all datasets, with 2-strobe mode accuracy being slightly higher. At the same time, all 3-strobes mode resulted in higher runtime than the 2-strobe mode, with some exceptions on the fruitfly dataset (Figure S16). Given that the 3-strobe modes added runtime overhead over the 2-strobe mode without demonstrating any accuracy benefits, only the 2-strobe multi-context seeds are used in strobealign v.0.17.0. We note that strobealign's mapping approach (syncmer thinned strobemers, chaining, and scoring and cutoffs) was designed around a two strobe approach. While a 3-strobe approach did not demonstrate benefit in strobealign, we don't exclude the possibility of an approach with three strobes working in a different application or context. It is also possible that 8 bits each (only 256 unique values) for the second strobe are too small and causing significant hash collisions on the larger datasets, which could explain the significant runtime increase (Figure S16). Additional implementation specific runtime optimizations for 3-strobes could also be made such as storing previously computed hashes during multi-context-seed construction, but we did not consider this given that we did not see substantial gain in accuracy.

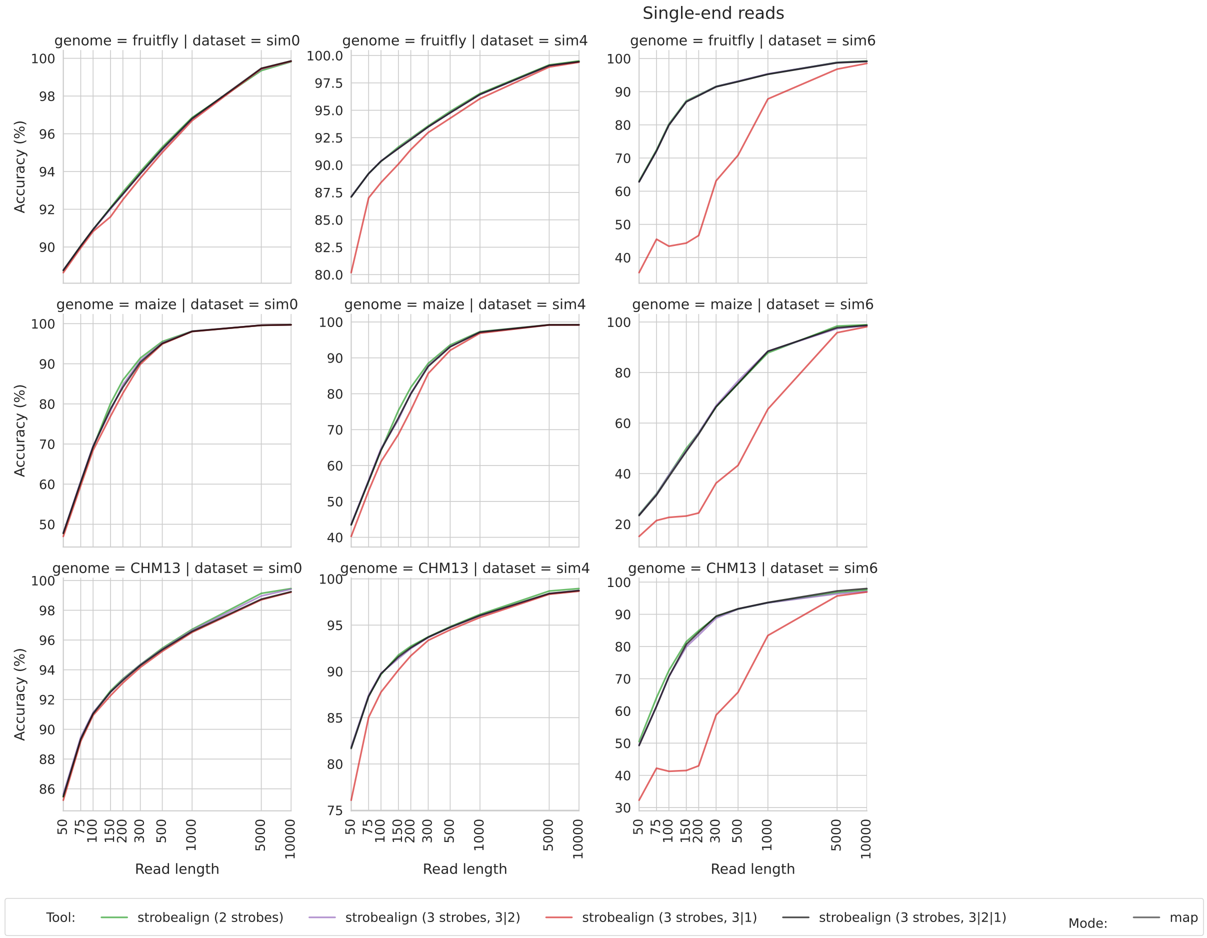


Fig. S15: Accuracy of strobealign in 3-strobe and 2-strobe modes, in mapping-only mode.

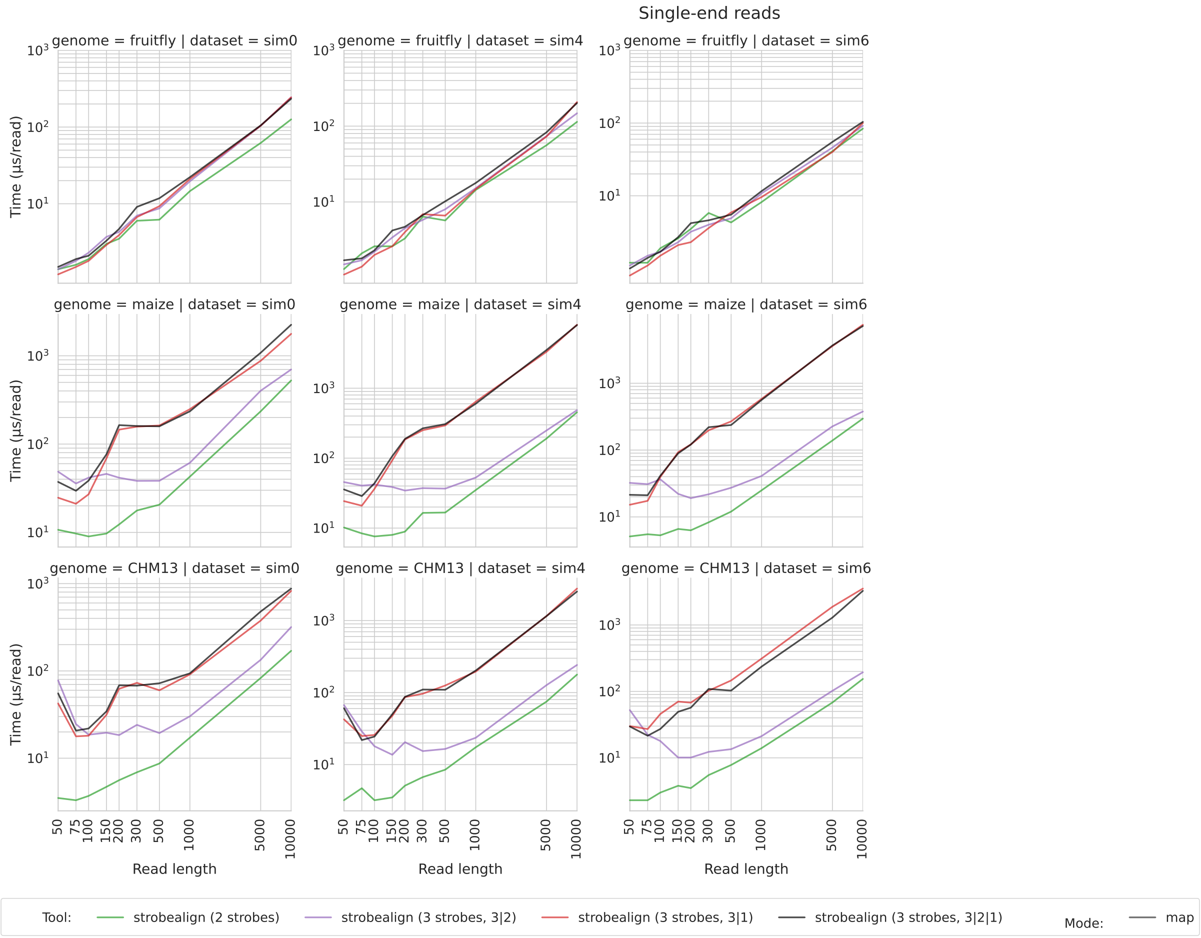


Fig. S16: Runtime of strobealign in 3-strobe and 2-strobe modes, in mapping only mode.

### 8 SNV and indel calling

We ran the SNV and indel calling benchmark as performed in strobealign [7] using bcftools as the variant caller and two simulated (SIM150, SIM250) and two biological (BIO150, BIO250) paired-end Illumina datasets. See [7] for details.

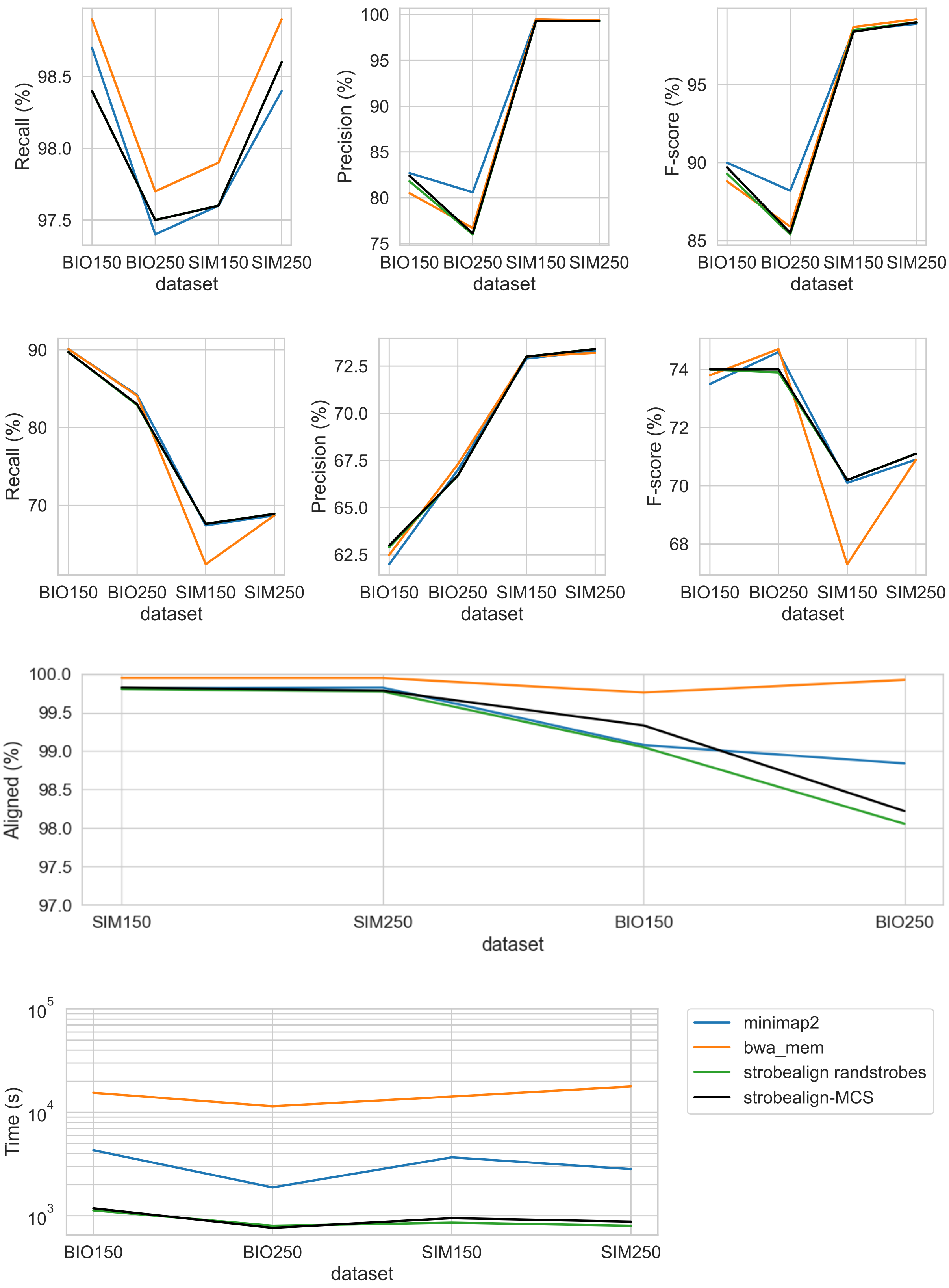


Fig. S17: Recall, precision and F-score for SIM150, SIM250, BIO150, BIO250. Top row shows SNV calling, second row from the top shows indel calling, third row shows the mapping rate, and bottom row runtime.

### 9 Chaining

We implemented a chaining algorithm largely inspired by the approach in minimap2 [4]. The goal of chaining (sometimes also referred to as *colinear chaining*) is to connect seed matches, called anchors, into longer chains that respect the same order in query and reference coordinates. Unlike minimap2, which operates on minimizer seeds, strobealign uses strobemers composed of two syncmers to construct chains. We split each strobemer into its constituent syncmers and treat each as an individual anchor $(x, y)$, where $x$ is the reference position and $y$ is the query position.

Given two anchors $a_j = (x_j, y_j)$ and $a_i = (x_i, y_i)$ with $x_j < x_i$ and $y_j < y_i$, we define

$$d_r = x_i - x_j, \qquad d_q = y_i - y_j.$$

The chaining score of extending $a_j$ to $a_i$ is defined as

$$s(i,j) \;=\; \min\{k, \min(d_q, d_r)\} \;-\; \big(\alpha \cdot |d_r - d_q| + \beta \cdot \min(d_q, d_r) + 0.5 \cdot \log_2(|d_r - d_q| + 1)\big),$$

where $k$ is the seed length and $\alpha, \beta$ are tunable parameters. The penalties account for gaps and diagonal deviations. The logarithmic penalty favors long indels over many short ones, reflecting the distribution of sequencing errors and biological variants. The above defined score function is used in minimap2 [4].

Anchors are sorted by reference, then query position. For each anchor $a_i$, the best score $S[i]$ of any chain ending at $i$ is

$$S[i] = \max_{j<i} \; S[j] + s(i,j),$$

restricted to predecessors within a fixed look-back window $h$ (by default $h = 50$ as in minimap2). This reduces runtime from $O(N^2)$ to $O(N \cdot h)$ for $N$ anchors. Additionally, anchors separated by more than a maximum reference gap $g_{\max}$ are skipped. These heuristics, the first taken from minimap2, keep the algorithm efficient for long noisy reads while preserving near-optimal chains in practice.

Chains are recovered by backtracing through predecessor pointers. We retain not only the best chain but also alternative chains above a relative score threshold, ensuring that repetitive or ambiguous mappings are preserved for downstream alignment. To avoid redundancy, anchors already used in higher-scoring chains are skipped when reconstructing additional chains. The resulting chains define candidate alignment intervals, which are then passed to the extension step.

[8] Ivan Tolstoganov, Marcel Martin, and Kristoffer Sahlin. Multi-context seeds enable fast and high-accuracy read mapping. *bioRxiv*, 2025.
